# USP9X deubiquitylates DVL2 to regulate WNT pathway specification

**DOI:** 10.1101/289595

**Authors:** Casey P. Nielsen, Kristin K. Jernigan, Nicole L. Diggins, Donna J. Webb, Jason A. MacGurn

## Abstract

The WNT signaling network is comprised of multiple receptors that relay various input signals via distinct transduction pathways to execute multiple complex and context-specific output processes. Integrity of the WNT signaling network relies on proper specification between canonical and non-canonical pathways, which presents a regulatory challenge given that several signal transducing elements are shared between pathways. Here, we report that USP9X, a deubiquitylase, and WWP1, an E3 ubiquitin ligase, regulate a ubiquitin rheostat on DVL2, a WNT signaling protein. Our findings indicate that USP9X-mediated deubiquitylation of DVL2 is required for canonical WNT activation, while increased DVL2 ubiquitylation is associated with localization to actin-rich projections and activation of the planar cell polarity (PCP) pathway. We propose that a WWP1-USP9X axis regulates a ubiquitin rheostat on DVL2 that specifies its participation in either canonical WNT or WNT-PCP pathways. These findings have important implications for therapeutic targeting of USP9X in human cancer.

## Introduction

The canonical WNT β-catenin signaling pathway is involved in regulating many cellular processes such as cell fate determination during embryonic development, cell proliferation, and adult tissue homeostasis. Thus, it is not surprising that aberrant activation of the canonical Wnt pathway is known to occur in many types of cancer (MacDonald et al., 2009; Saito-Diaz et al., 2013). There are also several noncanonical WNT signaling pathways including the WNT-Planar Cell Polarity (WNT-PCP) pathway which controls cell migration and tissue polarity. Dysregulation of the WNT-PCP pathway has been linked to cancer invasion and metastasis (Katoh, 2005; Luga et al., 2012; Wang, 2009). While the canonical WNT β-catenin pathway and the noncanonical WNT-PCP pathway use divergent effector mechanisms to regulate distinct cellular functions, these pathways share membrane receptor components and the cytoplasmic WNT transducer protein dishevelled (DVL). Despite its key role in both pathways, the mechanisms dictating DVL participation in canonical or noncanonical WNT signaling are yet to be elucidated.

Initiation of the canonical WNT β-catenin pathway occurs when extracellular WNT ligand binds to the co-receptors Frizzled (FZD) and low-density lipoprotein receptor-related protein 5/6 (LRP5/6) leading to recruitment of DVL and AXIN to the WNT ligand receptor complex (MacDonald et al., 2009). This ultimately results in the inhibition of β-catenin ubiquitylation and degradation such that stabilized β-catenin can enter the nucleus to initiate a transcriptional program (MacDonald et al., 2009; Saito-Diaz et al., 2013). On the other hand, core WNT-PCP pathway components Van Gogh-Like 1 (VANGL1), FZD, Prickle (Pk), DVL, and others function to activate RHOA, c-Jun N-terminal kinase (JNK), and nemo-like kinase (NLK) signaling cascades in order to coordinate tissue polarity and cell motility through regulation of actin dynamics (Glinka et al., 2011).

Ubiquitylation is known to be involved in key regulatory steps of both the canonical WNT and noncanonical WNT-PCP pathways. For example, ubiquitin-mediated regulation of cytoplasmic β-catenin stability is well characterized (Marikawa and Elinson, 1998). In addition, other steps of the WNT pathway upstream of β-catenin stabilization undergo regulation by the ubiquitin system. Notably, several members of the NEDD4 family of E3 ubiquitin ligases (SMURF1, ITCH, and NEDD4L) have been found to negatively regulate stability of WNT pathway components. SMURF1 interacts with and ubiquitylates AXIN, inhibiting its interaction with the WNT co-receptor LRP5/6 (Ding et al., 2013; Fei et al., 2014; Fei et al., 2013; Tanksley et al., 2013; Wei et al., 2012). Both ITCH and NEDD4L promote degradation of DVL2 (Cadavid et al., 2000; Ding et al., 2013; Fei et al., 2014; Fei et al., 2013; Wei et al., 2012). Additionally, the deubiquitylase (DUB) USP34 was found to antagonize ubiquitylation of AXIN, promoting its stabilization and function in the canonical WNT pathway (Lui et al., 2011). SMURF1 and SMURF2 negatively regulate the WNT-PCP pathway by targeting WNT-PCP receptor component Prickle1 for degradation (Narimatsu et al., 2009). Furthermore, DVL2 is also known to undergo positive regulation by the ubiquitin system. For example, K63-linked ubiquitylation of the N-terminal DAX domain, which is known to mediate dynamic polymerization of DVL2, has been implicated as a positive regulator of DVL2 signal transduction (Schwarz-Romond et al., 2007; Tauriello et al., 2010). These examples highlight the complex regulation of canonical and non-canonical WNT pathways by ubiquitin conjugation and deconjugation machinery.

The NEDD4 family of E3 ubiquitin ligases is conserved from yeast to humans and has been implicated in the ubiquitin-mediated endocytosis of many plasma membrane proteins including surface receptors, ion channels, and nutrient transporters (David et al., 2013; He et al., 2008; Hryciw et al., 2004; Kuratomi et al., 2005; Lee et al., 2011; Lin et al., 2011; MacGurn et al., 2012). There are 9 NEDD4 family E3 ubiquitin ligases encoded in the human genome, each with distinct physiological functions (Rotin and Kumar, 2009). While not all NEDD4 family E3 ubiquitin ligases have been characterized extensively, several have a strong preference for catalyzing K63-linked polyubiquitin chains *in vitro (Ingham et al., 2004; Kim and Huibregtse, 2009)*. Given the broad role of NEDD4 family E3 ubiquitin ligases in the regulation of membrane trafficking and cell signaling, it is not surprising that they are often found to be dysregulated in human diseases including cancer (Bernassola et al., 2008). NEDD4 family members have a conserved domain structure including an N-terminal C2 domain that binds lipids, a C-terminal HECT ubiquitin ligase domain, and 2-4 internal WW domains. The WW domains have a high affinity for PY (Pro-Pro-x-Tyr) motifs and mediate interactions with substrates, adaptors, and regulatory factors (Ingham et al., 2004). Here, we report that the NEDD4 family member WWP1 interacts with USP9X and DVL2, and we find that these associations are governed by interactions between the WW domains of WWP1 and PY motifs present in both USP9X and DVL2. Importantly, we find that these interactions control a ubiquitylation rheostat on DVL2 that regulates both canonical WNT and non-canonical WNT-PCP activation in human breast cancer cells. Thus, we find that this USP9X-DVL2-WWP1 axis is a key regulatory node in the WNT pathway with the potential to influence both canonical and noncanonical WNT signaling in the context of human cancer.

## Results

### The WW domains of WWP1 coordinate interactions with USP9X and DVL2

In an attempt to identify factors that regulate WWP1 function we used SILAC-based quantitative proteomics to generate a WWP1 interaction profile in MDA-MB-231 cells (a triple negative breast cancer cell line). This experiment identified interactions with various plasma membrane (PM) and PM-associated proteins, factors associated with membrane trafficking, signaling factors, and elements of the ubiquitin-proteasome system (**Table S1** and **FIG S1A**). To validate results obtained by quantitative proteomics, we tested for co-purification of a subset of putative WWP1 interacting proteins with FLAG-WWP1 stably expressed in MDA-MB-231 cells (**FIG 1A**). This approach confirmed FLAG-WWP1 interactions with several putative interactors – including USP9X, DVL2 and SPG20 – although some putative interactors such as ARRDC1 and USP24 could not be confirmed by this approach. Considering the possibility that some putative interacting proteins may artificially result from stable overexpression of FLAG-WWP1, we performed native co-immunoprecipitation of WWP1 and identified interactions with USP9X and DVL2 at endogenous expression levels (**FIG S1B**). Importantly, other studies have reported interactions between USP9X and NEDD4 family E3 ubiquitin ligases – including SMURF1 (Xie et al., 2013) and ITCH (Mouchantaf et al., 2006). Similarly, DVL2 was previously reported to interact with other NEDD4 family E3 ubiquitin ligases – including WWP2 (Mund et al., 2015) and NEDD4L (Ding et al., 2013; Wei et al., 2012). Notably, both USP9X and DVL2 contain multiple PY motifs capable of binding the WW domains of NEDD4 family E3 ubiquitin ligases (**FIG 1B**).

**Figure 1.**
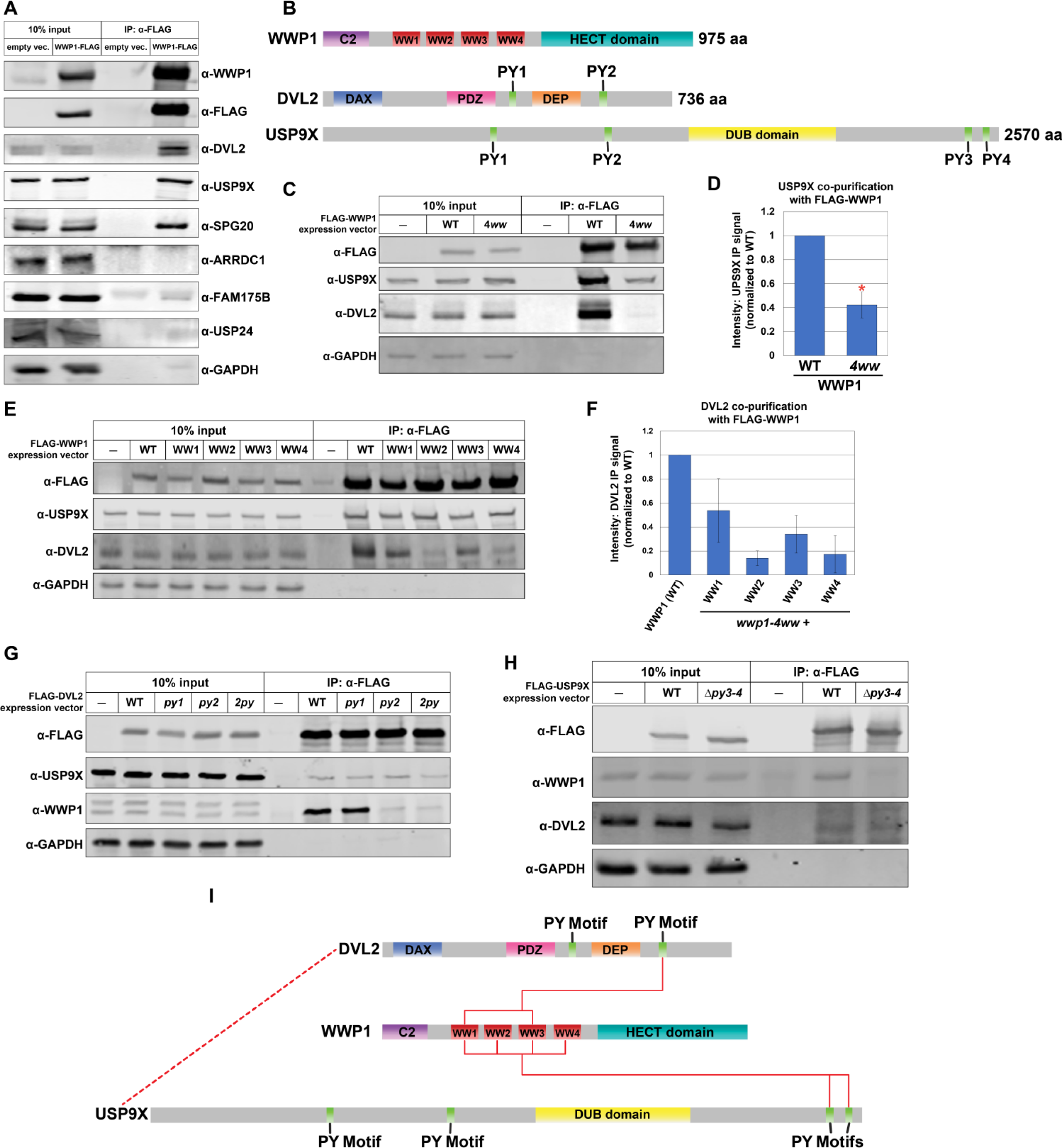
WWP1 interacts with USP9X and DVL2 in MDA-MB-231 human breast cancer cells. (**A**) Co-immunoprecipitation analysis testing hits from SILAC-MS analysis in (FIG S1A). FLAG-WWP1 was affinity purified from MDA-MB-231 cell lysates. Input and immunoprecipitated (IP) fractions were blotted for the indicated species. **(B)** Schematic of known domains in WWP1, DVL2, and USP9X. **(C)** Analysis of co-purification of DVL2 and USP9X with WWP1 variants. The indicated WWP1 variants were FLAG affinity purified, and quantitative immunoblots were performed to assess co-purification of interacting factors. The *wwp1-4ww* mutant indicates that all four WW domains have been mutated to disrupt the ability to bind PY motifs. **(D)** Quantification of USP9X co-purification with FLAG-WWP1 variants (normalized to WT) over multiple experiments (n=3). Red asterisk indicates a significant difference based on Student’s t-test analysis. **(E)** Analysis of co-purification of DVL2 and USP9X with WWP1 variants. The indicated WWP1 variants were FLAG affinity purified, and quantitative immunoblots were performed to assess co-purification of interacting factors. Each mutant variant of WWP1 contains a single in-tact WW domain (indicated), while the remaining three are mutated to disrupt PY motif binding. **(F)** Quantification of DVL2 co-purification with FLAG-WWP1 variants (normalized to WT) over multiple experiments (n=3). **(G)** Analysis of co-purification of WWP1 and USP9X with DVL2 variants. The indicated mutant DVL2 variants were FLAG affinity purified, and quantitative immunoblots were performed to assess co-purification of interacting factors. Mutant variants of DVL2 included point mutations disrupting the ability of its PY motifs to interact with WW domains. **(H)** Analysis of co-purification of WWP1 and DVL2 with USP9X variants. The indicated USP9X variants were FLAG affinity purified, and quantitative immunoblots were performed to assess co-purification of interacting factors. Mutant variants of USP9X included a small C-terminal truncation which results in deletion of both C-terminal PY motifs. **(I)** A schematic model for how USP9x and DVL2 both interact with WWP1 via PY motif interactions. The data also suggest an interaction between USP9X and DVL2 that is independent of WWP1 (dotted red line).

Based on our findings and previous reports we decided to further investigate the basis of physical interactions between WWP1, USP9X, and DVL2. Since USP9X and DVL2 both contain multiple PY motifs we hypothesized that the WW domains of WWP1 might function to scaffold these proteins into a complex. To test this, we generated WWP1 WW domain point mutant variants designed to disrupt PY motif interactions (WxxP → FxxA) (Chen et al., 1997; Ermekova et al., 1997; Gajewska et al., 2001) and analyzed DVL2 and USP9X interaction by co-immunoprecipitation. We found that disruption of individual WW domains within WWP1 did not diminish interaction with USP9X or DVL2 (**FIG S1C** and **S1D**). Indeed, we unexpectedly observed that the *wwp1-ww2* mutant exhibited significantly greater interaction with DVL2 (**FIG S1C** and **S1D**). Although individual WW domains were dispensable for interaction with DVL2 and USP9X, we found that a WWP1 variant with point mutations disrupting each WW domain (*wwp1-4ww*) exhibited significantly decreased interaction with USP9X and complete loss of interaction with DVL2 (**FIG 1C** and **1D**). Together, these findings indicate that ***(i)*** WW domain scaffolding partially contributes to the interaction between WWP1 and USP9X, ***(ii)*** WW domain interactions are required for WWP1 engagement with DVL2, and ***(iii)*** the WW domains of WWP1 function redundantly for interaction with both DVL2 and USP9X.

To further probe the WW domain specificity underpinning WWP1 interactions with DVL2 and USP9X, we generated a panel of WWP1 variants containing only a single in-tact WW domain (designated as *wwp1*-WW1, *wwp1*-WW2, *wwp1*-WW3, and *wwp1*-WW4). Importantly, we found that any individual WW domain is sufficient to restore interaction with USP9X, suggesting complete redundancy for this interaction (**FIG 1E**). In contrast, we found that only WW1 and WW3 were sufficient to restore interaction with DVL2 (**FIG 1E** and **1F**), indicating that WW2 and WW4 do not contribute to the interaction between WWP1 and DVL2. Furthermore, the observation that *wwp1*-WW2 and *wwp1*-WW4 interact with USP9X but not DVL2 indicates that the USP9X-WWP1 interaction occurs independent of DVL2 engagement. Overall, these findings suggest that the WW domains of WWP1 scaffold interactions with both USP9X and DVL2 in a manner that is redundant for USP9X and partially redundant for DVL2.

### PY Motifs in USP9X and DVL2 Promote Interaction with WWP1

Based on the observation that WW domains of WWP1 are important for interacting with USP9X and DVL2, we hypothesized that PY motifs in USP9X and DVL2 are required to engage WWP1. DVL2 contains two PY motifs (FPAY_393_ and PPPY_568_) that flank the DEP domain (**FIG 1B**). To test if either of these PY motifs are required for DVL2 to bind WWP1, we generated DVL2 variants with point mutations at individual PY motifs (FPAY_393_ → FAAA_393_ and PPPY_568_ → PAPA_568_) and performed co-immunoprecipitation analysis using FLAG-DVL2 as bait. This analysis revealed that PY1 (FPAY_393_) is dispensable for interaction with WWP1 while PY2 (PPPY_568_) is required to interact with WWP1 (**FIG 1G**). Furthermore, these results indicate that disruption of the DVL2-WWP1 interaction does not affect DVL2 engagement with USP9X (**FIG 1G**). Taken together, our results indicate that the DVL2-WWP1 interaction is governed by binding of the PY2 motif of DVL2 to the WW1 or WW3 domains of WWP1.

We hypothesized that PY motifs in USP9X also contribute to its interaction with WWP1. USP9X contains four PY motifs (QPQY_409_, HPRY_1017_, NPQY_2437_, and APLY_2515_), with the last two occurring at the C-terminus of the protein (**FIG 1B**). To test if these C-terminal PY motifs are required for interaction with WWP1, we generated a USP9X variant deleted for the C-terminal portion of the protein (USP9X_1-2433_, or *usp9x-Δpy3-4*). Importantly, deletion of the two C-terminal PY motifs resulted in complete loss of interaction with WWP1 (**FIG 1H**). However, point mutations in these C-terminal PY motifs (NPQY_2437_ → NAQA_2437_ (*py3*) and APLY_2515_ → AALA_2515_ (*py4*)) resulted in only partial loss of WWP1 interaction (**FIG S1E**). Notably, in each experiment loss of USP9X interaction with WWP1 did not affect its interaction with DVL2, indicating that the USP9X-DVL2 interaction occurs independently of WWP1. Thus, WWP1 engages both USP9X (partially) and DVL2 (primarily) via non-exclusive WW-PY interactions while DVL2 and USP9X exhibit the ability to interact independently of WWP1 (**FIG 1I**).

### WWP1 and USP9X establish a ubiquitin rheostat on DVL2

DVL2 was previously reported to be a substrate of several NEDD4 family E3 ubiquitin ligases including ITCH, NEDD4L, and NEDL1 (Ding et al., 2013; Miyazaki et al., 2004; Narimatsu et al., 2009; Nethe et al., 2012; Wei et al., 2012) although both positive and negative regulation of DVL2 by NEDD4 family members has been reported. However, WWP1 has not previously been reported to regulate DVL2. Given the physical associations between WWP1 and DVL2 we hypothesized that DVL2 might be subject to ubiquitin conjugation by the E3 activity of WWP1. We found that recombinant WWP1 exhibited ubiquitin conjugation activity toward HA-tagged DVL2 (purified from human STF-293 cell lysates) (**FIG 2A** and **FIG S2A-B**) and this activity is lost in a lysine-deficient variant of DVL2 (*dvl2-K0*, **FIG S2C**). Importantly, we also found that USP9X can reverse the ubiquitylation of DVL2 by WWP1 (**FIG 2B**), indicating that DVL2 is a substrate for both WWP1 and USP9X. Strikingly, the K48-specific deubiquitylase OTUB1 and the K63-specific deubiquitylase AMSH did not exhibit activity towards WWP1-ubiquitylated DVL2 (**FIG 2B**), suggesting that WWP1-conjugated ubiquitin on DVL2 is unlikely to contain K63- or K48-linked polymers.

**Figure 2.**
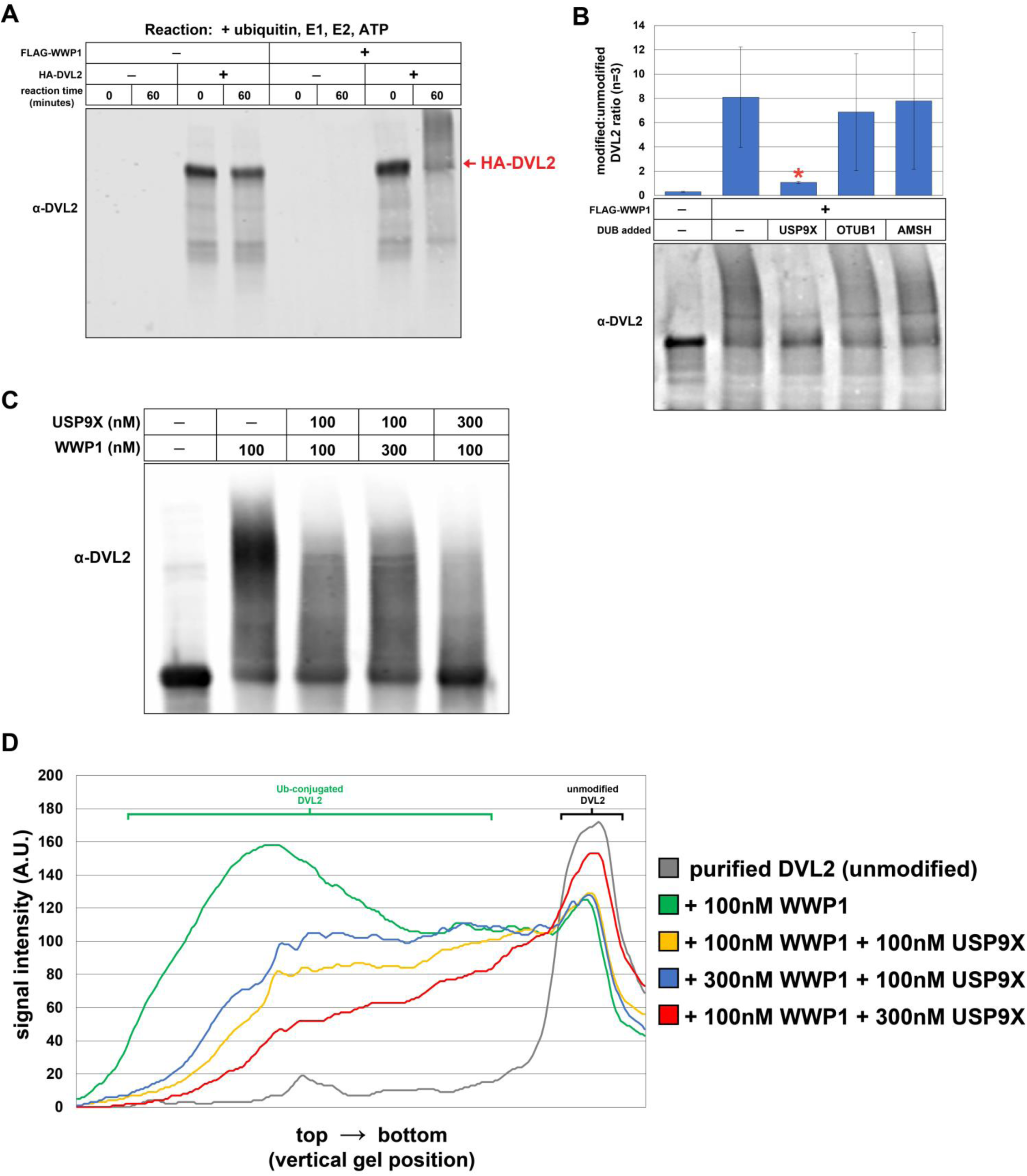
WWP1 and USP9X operate on DVL2 to establish a ubiquitylation rheostat. **(A)** *In vitro* ubiquitylation reactions were performed using recombinant purified FLAG-WWP1 (60nM) and HA-DVL2 purified from cultured cells. E3 conjugation reactions were allowed to proceed for 60 minutes before the conjugation reaction was terminated. **(B)** Deconjugation assays were performed using HA-DVL2 ubiquitylated *in vitro* by WWP1 (as generated in **(A)**). Equivalent molar ratios of USP9X, OTUB1 and AMSH (100nM) were added and each reaction was allowed to proceed for 60 minutes before the deconjugation reaction was terminated and HA-DVL2 was resolved by SDS-PAGE and immunoblot (bottom panel). For each sample, a ratio of modified:unmodified HA-DVL2 was measured using ImageJ. The modified:unmodified HA-DVL2 ratio was averaged over multiple replicate experiments (n=3). The red asterisk indicates a statistically significant difference (p<0.05) compared to DVL2 that has been modified by WWP1 and subsequently treated with OTUB1 or AMSH. **(C)** Representative immunoblot of *in vitro* ubiquitylation/deubiquitylation reactions (replicate experiments shown in **FIG S2D-E**). Experiments were performed using indicated combinations of purified FLAG-WWP1 and 6xHIS-USP9X with HA-DVL2 purified from cultured cells (see **FIG S2A-B**). Reactions were allowed to proceed for 60 minutes before being terminated and HA-DVL2 was resolved by SDS-PAGE and immunoblot. **(D)** Line density plots from the HA-DVL2 immunoblot data in **(C)** illustrate differences in extent of ubiquitylation in each sample. Replicate experiments are shown in **FIG S2D-E**.

Since WWP1 and USP9X both interact with and operate on DVL2, we hypothesized that the coordination of WWP1 E3 ubiquitin ligase activity and USP9X deubiquitylase activity might serve to establish a ubiquitin rheostat on DVL2. To test this, we incubated DVL2 with different molar ratios of WWP1 and USP9X and analyzed the outcome. Importantly, we found that different molar ratios of WWP1 and USP9X affected the extent of ubiquitylation on DVL2 (**FIG 2C-D**, **S2D** and **S2E**). These findings indicate that coordinated WWP1 and USP9X activities can establish a ubiquitin rheostat on DVL2.

### USP9X promotes canonical WNT activation

Given the established function of DVL2 as a key signal transducing protein in the WNT pathway and given that NEDD4 family members have been reported to negatively regulate WNT signaling (Tanksley et al., 2013) potentially via ubiquitylation of DVL2 (Ding et al., 2013), we decided to test if USP9X regulates WNT signaling. We found that knockdown or knockout of *usp9x* attenuated WNT activation as measured by a TopFLASH reporter assay (**FIG 3A** and **S3A**) and by β–catenin stabilization (**FIG 3A**). The WNT signaling defect observed in *usp9x* knockout cells was complemented by transient expression of wildtype FLAG-USP9X but not a catalytic dead (C1556V, or CD) variant (**FIG 3B**), indicating that catalytic activity of USP9X is required for canonical WNT activation. Furthermore, we found that treating cells with WP1130, a small molecule known to inhibit the deubiquitylase activity of USP9X (Kapuria et al., 2010), inhibited canonical WNT activation by WNT3a with an IC_50_ of 2.9µM (**FIG 3C**). This result is consistent with our findings that USP9X is required for canonical WNT activation, however WP1130 is known to target other deubiquitylases (USP24, USP5, USP14, and UCH37) (de Las Pozas et al., 2018; Kapuria et al., 2010; Kushwaha et al., 2015; Peterson et al., 2015). Thus, we cannot exclude the possibility that inhibition of multiple DUB activities drives loss of WNT activation in the presence of WP1130. To further explore the possibility that USP9X activity promotes WNT activation we transiently transfected expression vectors for FLAG-USP9X (wildtype or catalytic dead) and measured WNT activation in the context of WNT3a stimulation. Importantly, we found that transient expression of wildtype FLAG-USP9X, but not a catalytic-dead variant, resulted in increased activation of the canonical WNT pathway (**FIG 3D**).

**Figure 3.**
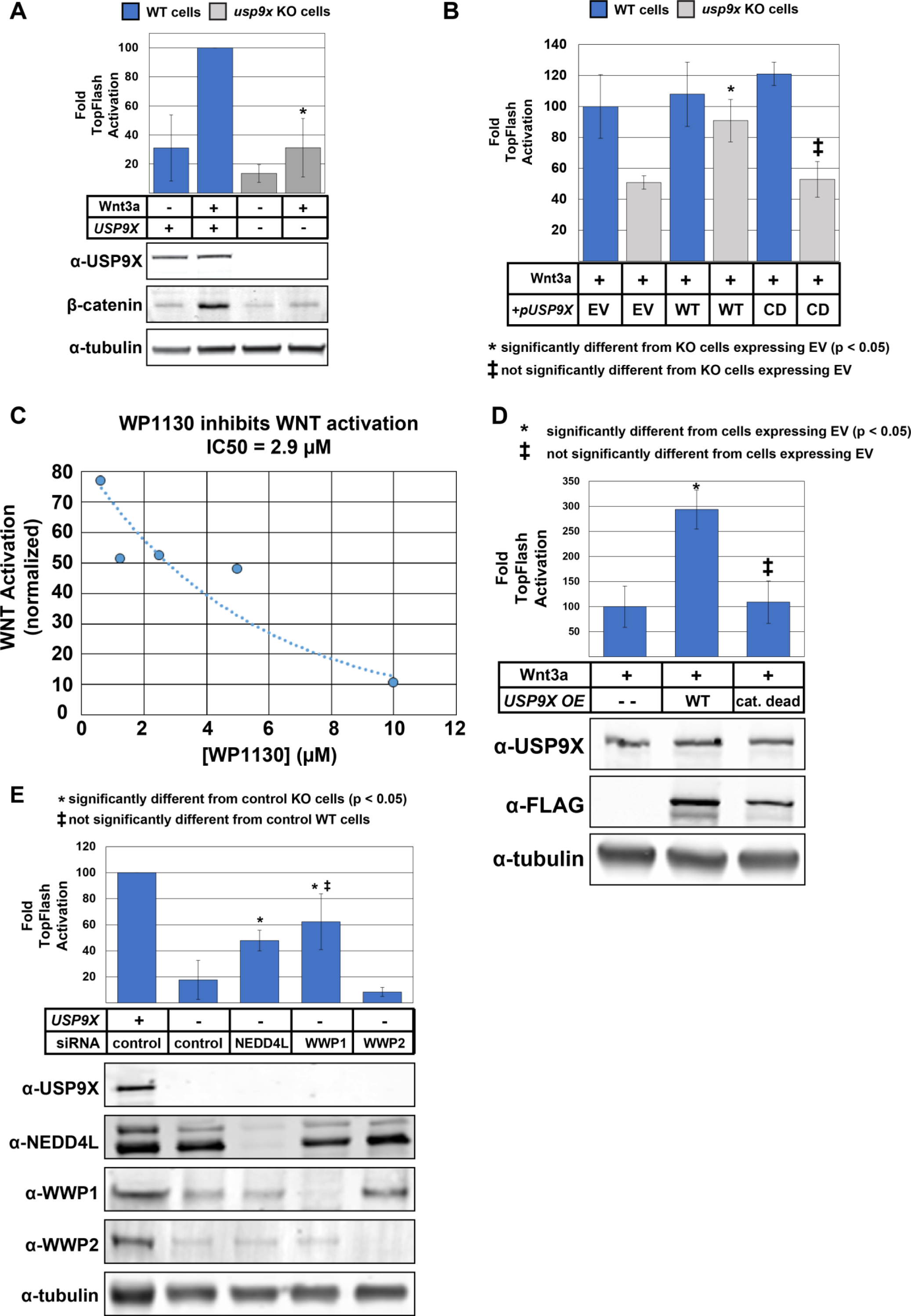
USP9X promotes canonical WNT activation. **(A)** Analysis of ligand-stimulated WNT activation in MDA-MB-231 cells (blue bars) and *usp9x* knockout equivalents (gray bars). TopFLASH luciferase assays were used to measure WNT activation (top panel) and immunoblotting was performed to assess stabilization of nuclear β-catenin. * indicates WNT activation in *usp9x* knockout cells is significantly decreased (p<0.05) compared to sibling MDA-MB-231 cells. **(B)** MDA-MB-231 cells (wildtype and *usp9x* knockout) were stimulated with WNT3a ligand and R-spondin and WNT activation was measured by TopFLASH luciferase assays. Prior to activation, cells were transiently transfected with empty vector (EV) or vector expression USP9X (wildtype (WT) or catalytic dead (CD; C1566V)). **(C)** Ligand-stimulated WNT activation was measured in the presence of indicated concentrations of WP1130. Based on the data an IC50 was estimated. **(D)** MDA-MB-231 cells were transfected with empty vector or plasmids for CMV-driven expression of wildtype (WT) or catalytic dead USP9X. **(E)** Ligand-stimulated WNT activation was measured using TopFLASH luciferase assays for MDA-MB-231 cells and *usp9x* knockout equivalents. Cells were transfected with control siRNA or siRNA targeting knockdown of NEDD4L, WWP1 or WWP2.

We hypothesized that loss of canonical WNT activation in the absence of USP9X may be due to unchecked activity of WWP1 or other NEDD4 family E3 ubiquitin ligases. Importantly, we found that the loss of canonical WNT activation in *usp9x* knockout cells was suppressed by coordinate knockdown of WWP1 or NEDD4L but not WWP2 (**FIG 3E**) suggesting that hyper-activation of multiple NEDD4 family members, including NEDD4L and WWP1, may attenuate WNT signaling in the absence of USP9X. Furthermore, we found that transient overexpression of wildtype WWP1, but not a catalytic dead variant (C890S), inhibited TopFLASH activation by WNT3a (**FIG S3B** and **S3C**). Unexpectedly, we also found that expression of the *wwp1-ww2* variant (harboring point mutations in WW2) inhibited canonical WNT activation to a greater extent that wildtype WWP1 (**FIG S3C**) while expression of the *wwp1-*WW2 variant (where all WW domains are disrupted except for WW2) exhibited hyperactivation of canonical WNT in the presence of WNT3a (**FIG S3B**). Thus, ability of WWP1 to antagonize canonical WNT activation correlates with its ability to bind DVL2 (**FIG 1E-F** and **S1C-D**). Taken together, these findings indicate that USP9X promotes canonical WNT activation while WWP1 antagonizes canonical WNT activation.

### USP9X regulation of WNT requires DVL2 deubiquitylation

Given our finding that WWP1 and USP9X interact with and operate on DVL2, we hypothesized that USP9X regulates canonical WNT activation by affecting the ubiquitylation status of DVL2. To address the mechanism of canonical WNT regulation by USP9X, we first tested if USP9X is required for β-catenin stabilization by LiCl – an inhibitor of GSK3. Importantly, LiCl-mediated stabilization did not require USP9X (**FIG S4A**), indicating USP9X functions upstream of the β-catenin destruction complex. Importantly, we found that knockdown or knockout of USP9X resulted in decreased steady state abundance of DVL2 protein and mobility shifts by SDS-PAGE consistent with hyper-ubiquitylation (**FIG 4A** and **S4B**). To identify post-translational modifications on DVL2 regulated by USP9X we performed a SILAC-MS on FLAG-DVL2 affinity purified from MDA-MB-231 cells and *usp9x* knockout equivalents. This analysis resolved many phospho-peptides and ubiquitin remnant (diGly) peptides derived from DVL2 with four internal lysines exhibiting elevated ubiquitylation in the absence of USP9X (**FIG 4B** and **FIG S4C-E**). Although ubiquitylation at N-terminal lysines of DVL2 has been reported (Madrzak et al., 2015; Tauriello et al., 2010) the ubiquitin modification events described in this study (K343, K428, K477 and K615) have not previously been reported.

**Figure 4.**
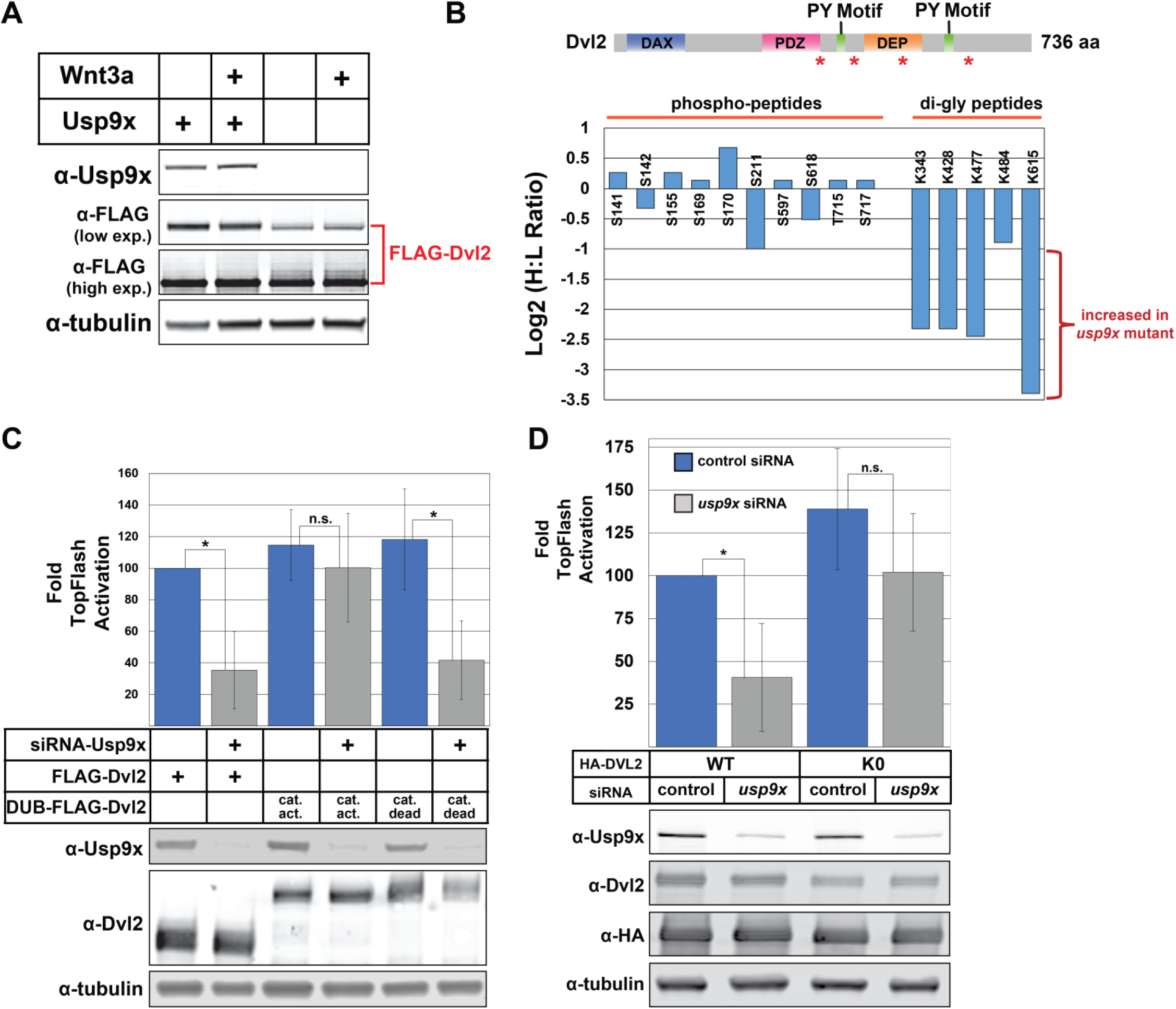
USP9X promotes canonical WNT activation via deubiquitylation of DVL2. **(A)** Immunoblotting analysis of FLAG-DVL2 mobility by SDS-PAGE from lysates of MDA-MB-231 cells or *usp9x* knockout equivalents. A low exposure α-FLAG blot is shown to illustrate differences in DVL2 abundance, while the higher exposure α-FLAG is shown to illustrate the higher MW species evident in *usp9X* knockout cells. **(B)** SILAC-MS profiling of post-translational modification sites (phosphorylation and ubiquitylation) on DVL2. In this experiment, FLAG-DVL2 purified from MDA-MB-231 cells (heavy) was compared to *usp9x* knockout equivalents (light). A schematic illustrating the domain structure of DVL2 is shown at the top with asterisks to indicate the position of detected ubiquitylation events. **(C)** Analysis of ligand-stimulated WNT activation was measured using TopFLASH reporter assays in STF-293 cells transfected with the indicated expression vectors. UL36 is the DUB domain fused to DVL2 in these experiments. **(D)** Following siRNA knockdown of USP9X, ligand-stimulated WNT activation was measured using TopFLASH reporter assays in STF-293 cells transfected with the indicated HA-DVL2 expression vectors. DVL2-K0 is a variant where all encoded Lys residues are substituted with Arg residues. * represents a statistically significant difference when compared to indicated sample (p<0.05). n.s. represents no significant difference relative to indicated comparison.

To test if USP9X-mediated regulation of WNT signaling requires DVL2 ubiquitylation we expressed DVL2 fused to the UL36 DUB domain (from human herpes virus) and found that this (but not a catalytic dead UL36 fusion) suppressed the WNT activation defect in the absence of USP9X (**FIG 4C**). These results indicate that USP9X regulates WNT signaling at (or in very close proximity to) the level of DVL2. To explore the possibility that USP9X-mediated deubiquitylation of DVL2 at specific sites is required for WNT activation we tested a panel of DVL2 lysine mutants (K343R, K429R, K477R and K615R) but found expression of these mutants was insufficient to suppress the WNT signaling defect observed in the absence of USP9X (**FIG S4F**). We next considered the possibility that USP9X may regulate multiple ubiquitylation events on DVL2, as suggested by SILAC-MS analysis (**FIG 4B**). Importantly, we found that expression of a “lysine-less” DVL2 variant with all encoded lysine residues switched to arginine (DVL2-K0) is sufficient to restore WNT activation in the absence of USP9X (**FIG 4D**). Taken together, these results indicate that increased ubiquitylation of DVL2 is required for the attenuation of WNT signaling observed in the absence of USP9X. Thus, the USP9X-mediated deubiquitylation of DVL2 is critical for activation of canonical WNT signaling.

### DVL2 ubiquitylation status regulates interactions with canonical and WNT-PCP factors

We hypothesized that the altered ubiquitylation status of DVL2 in the absence of USP9X regulates its function as a signal transduction protein. To test this hypothesis, we first characterized the DVL2 interaction network in MDA-MB-231 cells using SILAC-MS (**TABLE S2, TABLE S3**, and **FIG 5A**). These experiments revealed two near-stoichiometric interactions with the AP-2 clathrin adaptor complex and with the non-canonical WNT factors VANGL1 and cofilin which are known to participate in the planar cell polarity (PCP) pathway (Luga et al., 2012). Additionally, we identified several sub-stoichiometric interactions with NEDD4 family E3 Ub ligases and proteasome subunits (**TABLE S2, TABLE S3**, and **FIG 5A**). Many of these interactions had been previously reported (Angers et al., 2006; Yu et al., 2007). Several interactions were confirmed by co-IP (**FIG 5B**) and co-localization by immunofluorescence microscopy (**FIG S5A**). Consistent with previous reports (Yu et al., 2007; Yu et al., 2010) we also found that knockdown of AP-2 complex subunits inhibited canonical WNT activation in a TopFLASH assay (data not shown).

**Figure 5.**
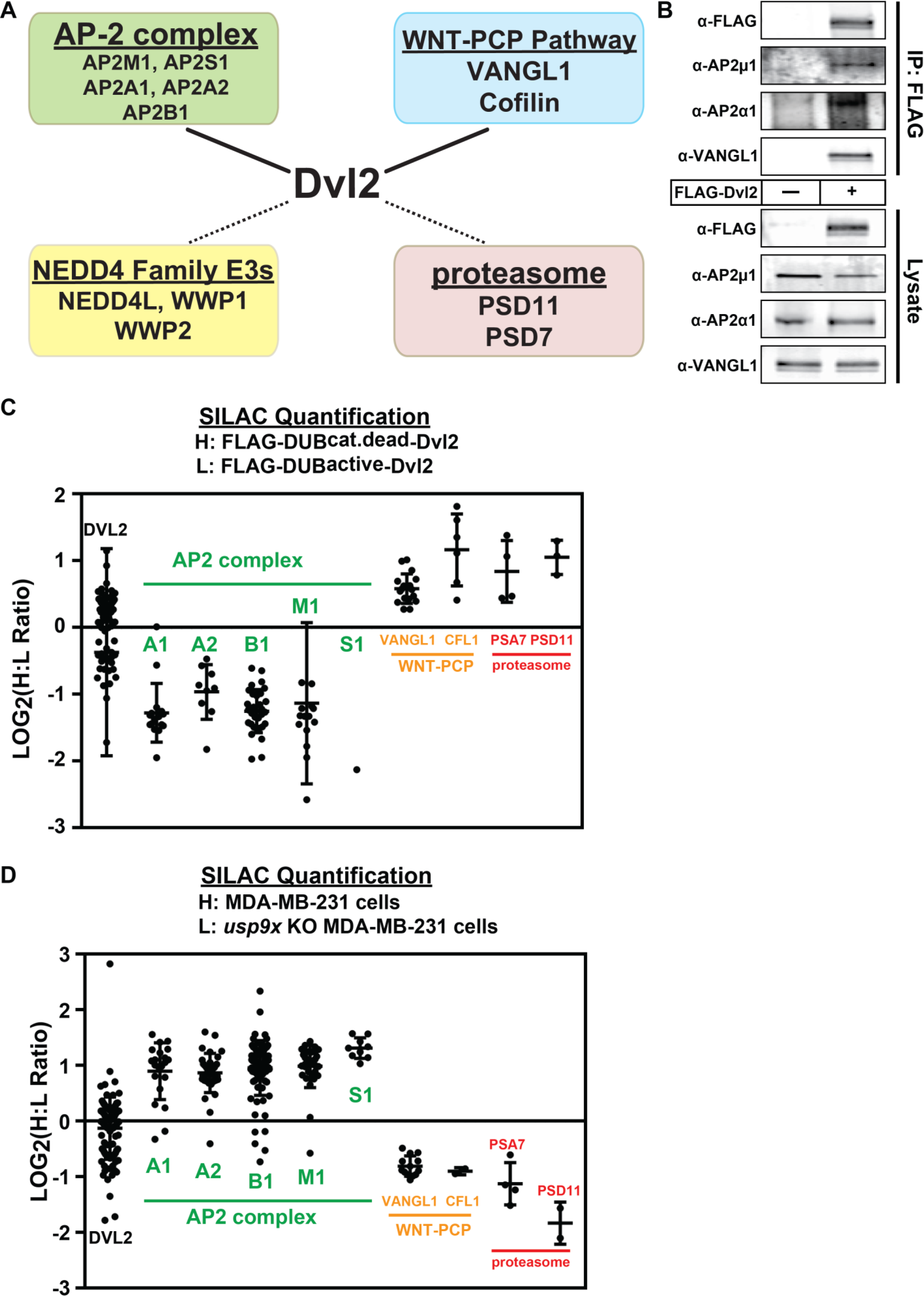
DVL2 ubiquitylation state specifies association with canonical WNT or WNT-PCP pathway factors. **(A)** SILAC-MS was used to resolve the DVL2 interaction network in MDA-MB-231 human breast cancer cells. Near-stoichiometric interactions (solid lines) were detected with the AP-2 clathrin adaptor complex and VANGL1-cofilin, which comprise a subcomplex of the WNT-PCP pathway. Sub-stoichiometric interactions (dotted lines) were identified for NEDD4 family E3 ubiquitin ligases and proteasome subunits. **(B)** Interactions detected by SILAC-MS were confirmed by coimmunoprecipitation. **(C and D)** SILAC-MS analysis was performed to measure how DVL2 interactions are impacted by catalytic active vs. catalytic dead DUB fusion **(C)** or by loss of *USP9X* **(D)**. Scatter plots depict individual H:L measurements (LOG_2_ transformed) for each peptide resolved for the indicated proteins. The mean value and standard deviation for each protein is indicated. For the experiment shown in **(C)**, several peptides with extremely low H:L ratios are not depicted here but are shown in **FIG S5B**. These outlier peptides correspond to sites of ubiquitin modification (**FIG S5C**) shown in **FIG 4B**. Differences between DVL2 (bait) and all interacting proteins (AP2 complex, VANGL1, CFL1, and proteasome subunits) are statistically significant (p<0.005).

To quantify how DVL2 interactions are affected by ubiquitylation, we performed SILAC-MS proteomic analysis comparing the interaction profile of catalytic active (light) or catalytic dead (heavy) UL36-DVL2 fusions (also used in **FIG 4C**). This revealed that catalytic dead UL36-DVL2 fusion (which can be stably ubiquitylated) interacts more with proteasome subunits and VANGL1/cofilin while the catalytic active UL36-DVL2 fusion (which is not stably ubiquitylated) interacts more with the AP-2 complex (**FIG 5C**). Importantly, we noticed in this experiment a small set of peptides derived from the DVL2-UL36 catalytic active fusion protein and absent from the DVL2-UL36 catalytic dead fusion protein (**FIG S5B**). Interestingly, these outlier peptides all correspond to tryptic events that can only occur in the absence of ubiquitylation (at positions K428, K477 and K615) (**FIG S5C**) indicating that these sites are largely ubiquitin-modified in the DVL2-UL36 catalytic dead fusion protein. We also used SILAC-MS to quantify how DVL2 interactions are altered in *usp9x* knockout cells and found similar results: DVL2 interacted more with VANGL1 and cofilin in *usp9x* knockout cells (where DVL2 ubiquitylation is elevated) but interacted more with AP-2 in the presence of USP9X (where DVL2 is less ubiquitylated) (**FIG 5D**). These results indicate ubiquitylated DVL2 interacts more with VANGL1/cofilin – components of the WNT-PCP pathway – while deubiquitylated DVL2 interacts more with the AP-2 complex, which is critical for canonical WNT activation.

### USP9X regulates cellular distribution of DVL2 and antagonizes WNT-PCP

Previous studies have reported that VANGL1 and cofilin localize to F-actin rich projections in migrating MDA-MB-231 cells (Hatakeyama et al., 2014) and that DVL2 is required for WNT5a-induced cell migration in these cells (Zhu et al., 2012). Based on our findings that loss of USP9X results in increased ubiquitylation of DVL2 and increased interactions between DVL2 and VANGL1/cofilin we hypothesized that loss of USP9X may result in re-localization of DVL2 to actin-rich projections in MDA-MB-231 cells. To test this, we used immunofluorescence microscopy to characterize FLAG-DVL2 distribution in MDA-MB-231 cells in the presence or absence of USP9X. In MDA-MB-231 cells treated with control siRNAs, DVL2 exhibited a punctate distribution throughout the cell body, reflective of previous descriptions of DVL2 localization (**FIG 6A, top panels**) (Schwarz-Romond et al., 2005) whereas MDA-MB-231 siUSP9X knockdown cells displayed a redistribution of DVL2 to actin-rich projections (**FIG 6A, middle panels**). Quantification of the cellular distribution of DVL2 in the presence and absence of USP9X reveals a significant increase in DVL2 in actin-rich projections upon loss of USP9X (**FIG 6B**). Importantly, coordinate knockdown of WWP1 prevented this re-distribution of DVL2, indicating that WWP1 activity is required for re-distribution of DVL2 to actin-rich projections in the absence of USP9X (**FIG 6A, bottom panel** and **FIG 6B**). Thus, our data indicates that USP9X promotes DVL2 localization to puncta that also contain AP-2 (**FIG S5A**), while loss of USP9X activity results in WWP1-dependent re-distribution of DVL2 to actin-rich projections in MDA-MB-231 breast cancer cells.

**Figure 6.**
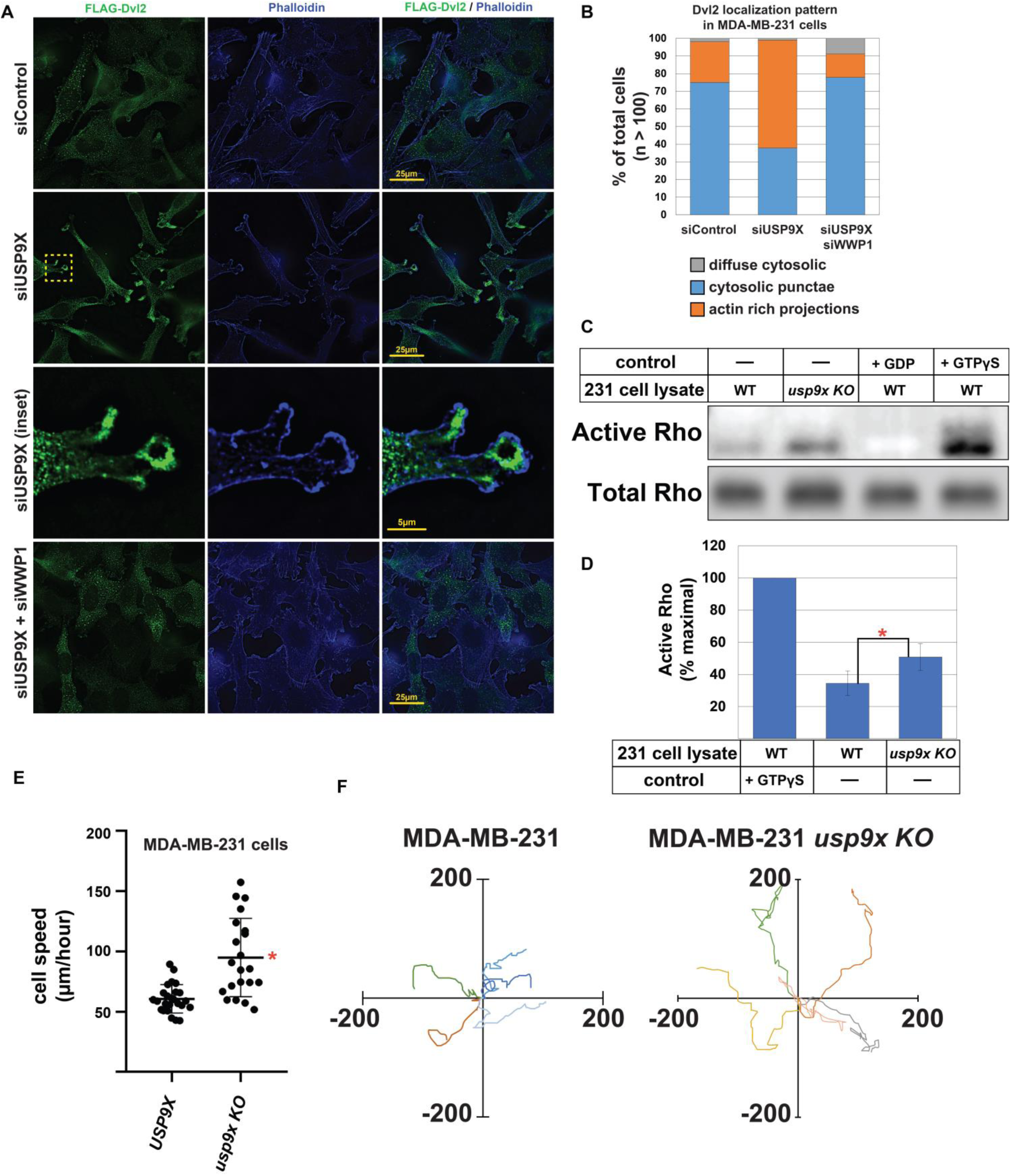
USP9X regulates DVL2 localization and antagonizes WNT-PCP activation. **(A)** Immunofluorescence imaging of FLAG-DVL2 (green) and actin (phalloidin, blue) in MDA-MB-231 human breast cancer cells transfected with control siRNA (top panels), siRNA targeting USP9X (middle panels), or a combination of siRNA targeting USP9X and siRNA targeting WWP1 (bottom panels). **(B)** Quantification of DVL2 cellular distribution over a population of MDA-MB-231 cells (n > 100). **(C)** WT MDA-MB-231 and *usp9x* KO variant cell lysates were assayed for active Rho by pulldown assay. Addition of GDP or non-hydrolyzable GTP (GTPƔS) to cell lysates served as negative and positive controls, respectively. **(D)** Quantification of Rho activation shown in **FIG 6C** (n=3) was performed by measuring the amount of active Rho detected in parent and *usp9x* knockout cells and normalizing to the GTPγS (maximal activation) control. The red asterisk indicates a significant difference when compared to WT cells (p<0.05) **(E)** MDA-MB-231 cells were plated at low density and cell migration (µm/hour) speed was measured. Red asterisk indicates a significant difference between the tested populations (n>30). **(F)** Rose plots showing trajectories of individual MDA-MB-231 cells (left) and *usp9x* knockout equivalents (right).

One feature of WNT5a-mediated PCP activation is Rho activation, which is DVL2-dependent and required for induction of cell migration by the WNT-PCP pathway (Zhu et al., 2012). Given our findings, we hypothesized that DVL2 re-distribution to actin-rich projections in MDA-MB-231 cells lacking USP9X might result in Rho activation. Indeed, we found that MDA-MB-231 cells normally exhibit about 35% of maximal Rho activation, while cells lacking USP9X exhibit 50% maximal Rho activation (**FIG 6C-D**). Taken together, these findings indicate that loss of USP9X in MDA-MB-231 cells results in re-distribution of DVL2 to actin-rich projections and Rho activation, which is similar to what occurs during WNT5a-mediated activation of the WNT-PCP pathway in these cells (Luga et al., 2012; Zhu et al., 2012).

### USP9X regulates cell motility in a manner dependent on DVL2 ubiquitylation

VANGL1, cofilin, and DVL2 are known to coordinate WNT-PCP activation to regulate actin dynamics and promote cell motility, migration and metastasis of MDA-MB-231 cells (Katoh, 2005; Luga et al., 2012; Yang and Mlodzik, 2015; Zhu et al., 2012). Based on these previous studies, we hypothesized that USP9X-mediated regulation of DVL2 may antagonize WNT-PCP-mediated cellular motility. Strikingly, we found that MDA-MB-231 breast cancer cells lacking USP9X exhibited a significant increase in motility compared to wildtype equivalents (**FIG 6E-F**). This increase in cell motility was complemented in cells transiently transfected with GFP-USP9X (**FIG S6A**). Consistent with this observation, we also found that treatment of MDA-MB-231 cells with WP1130 – a small molecule inhibitor of USP9X – also resulted in a significant increase in the speed of cell motility (**FIG S6B**). Importantly, the increased cell speed observed in *usp9x* knockout cells was dependent on VANGL1 (**FIG S6C**) indicating the increased motility phenotype requires and in-tact WNT-PCP pathway. To test if the increase cell motility observed in *usp9x* mutant cells was due to increased autocrine signaling, we treated cells with LGK974, a porcupine inhibitor that blocks the secretion of many WNT ligands, and found this had no effect on the speed of usp9x knockout MDA-MB-231 cells (**FIG S6D**). Taken together, these results indicate that loss of USP9X leads to aberrant activation of PCP-mediated motility independent of autocrine signaling.

We hypothesized that the elevated ubiquitylation of DVL2 might contribute to the increased motility phenotype observed in usp9x knockout MDA-MB-231 cells. Consistent with this hypothesis, we observed that knockdown of WWP1 suppressed the increased motility phenotype of *usp9x* knockout cells (**FIG 7A**), although transient overexpression of WWP1 did not alter speed of MDA-MB-231 cells (**FIG S7A**). Notably, MDA-MB-231 cells transiently expressing GFP-DVL2 did not exhibit any change in cell motility compared to untransfected control cells (**FIG S7B**), however expression of either wildtype or lysine-deficient (K0) GFP-DVL2 fully suppressed the increased motility phenotype of *usp9x* knockout MDA-MB-231 cells (**FIG S7C-D**). This result implicates DVL2 as a mediator of the cell motility phenotype observed for *usp9x* knockout cells, but it does not inform as to the role of ubiquitylation since both wildtype and lysine-deficient DVL2 fully suppress the *usp9x* knockout phenotype. To test the role of ubiquitylation, we analyzed cell motility of MDA-MB-231 cells stably expressing DVL2-UL36 (catalytic active and catalytic dead) DUB fusion proteins. Strikingly, we found that knockdown of *USP9X* resulted in increased motility of control cells and cells expressing catalytic dead DVL2-UL36 fusions, but not of cells expressing catalytic active DVL2-UL36 fusions (**FIG 7B).** This result reveals that the increased motility phenotype observed in the absence of *USP9X* requires increased ubiquitylation of DVL2. Finally, we tested if DVL2 ubiquitylation is required for WNT5a-induced cell motility in MDA-MB-231 cells. Interestingly, we found that expression of catalytic active DVL2-UL36 resulted in a significant dampening of WNT5a-induced cell motility compared to control cells expressing catalytic dead DVL2-UL36, although the observed inhibition was partial (**FIG 7C**). These results indicate that USP9X is a critical antagonist of PCP activation and cell motility in MDA-MB-231 cells and that this function is dependent on the ubiquitylation of DVL2. Based on our findings, we propose that the balance of USP9X and WWP1 activities toward DVL2 regulates WNT pathway specification, with USP9X-mediated DVL2 deubiquitylation promoting canonical WNT signaling and WWP1-mediated DVL2 ubiquitylation driving WNT-PCP activation and cell migration (**FIG 7D**).

**Figure 7.**
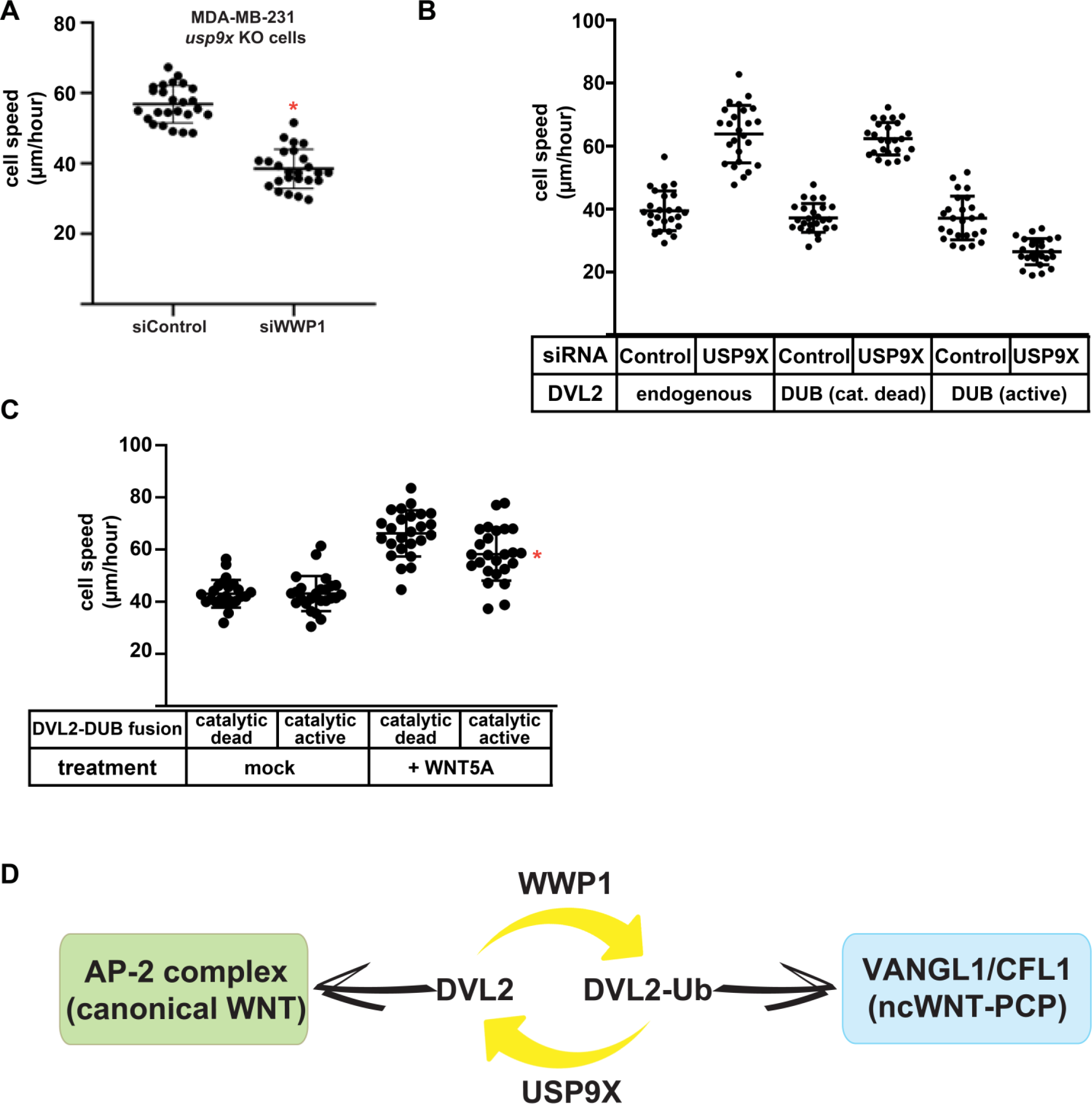
USP9X-mediated regulation of cell motility requires DVL2 ubiquitylation. **(A)** Migration speed of MDA-MB-231 *usp9x* KO cells treated with either control siRNA or siRNA targeting WWP1 was measured after plating at low density for the indicated conditions. Red asterisk indicates a significant difference compared to control siRNA treated cells (p<0.005). **(B)** MDA-MB-231 cell lines stably expressing DVL2-DUB fusions (catalytic active and catalytic dead) were treated with control siRNA or siRNA targeting USP9X and cell speed was measured for 25 cells. **(C)** MDA-MB-231 cell lines stably expressing DVL2-DUB fusions (catalytic active and catalytic dead) underwent mock or WNT5a treatment and cell speed was measured for 25 cells. The red asterisk indicates a statistically significant difference (p<0.005) compared to MDA-MB-231 cells stably expressing the catalytic dead DUB-DVL2 fusion treated with WNT5a. **(D)** Model for DVL2 regulation by the USP9X-WWP1 axis to establish a ubiquitin rheostat on DVL2 that influences interaction preference with either the clathrin adaptor AP-2 complex, which is important for canonical WNT signaling, or the VANGL1-cofilin complex, which is important for the WNT-PCP pathway. Thus, the USP9X-WWP1 axis regulates DVL2 ubiquitylation to determine WNT pathway specification.

## Discussion

Dishevelled has long been known to play a key role in the transduction of WNT receptor signaling, yet how it is regulated to specify participation in either canonical or non-canonical relays is not well understood. Here, we report that the NEDD4 family E3 ubiquitin ligase WWP1 interacts with DVL2 and the deubiquitylase USP9X, and that these interactions govern a ubiquitylation rheostat on DVL2 critical for both canonical WNT and WNT-PCP activation. Specifically, our data indicates that ***(i)*** USP9X, WWP1 and DVL2 physically associate to establish a ubiquitin rheostat on DVL2, ***(ii)*** USP9X-mediated deubiquitylation of DVL2 is required for canonical WNT activation, and ***(iii)*** USP9X antagonizes activation of the WNT-PCP pathway (and thus cell motility) by limiting the accumulation of ubiquitylated DVL2. Based on the data, we propose that the USP9X-WWP1 axis biases DVL2 engagement with either the canonical WNT pathway or the noncanonical WNT-PCP pathway (**FIG 7D**).

### A Novel Regulatory Axis for WNT Pathway Specification

The functional diversity of WNT signaling is a result of the complex network design that inter-weaves canonical and non-canonical elements to coordinate multiple specific signaling outputs. Thus, it is important to understand the network design features that couple canonical and non-canonical pathways and contribute to pathway specificity. The role of USP9X in WNT pathway specification draws parallels to diversin, an ankyrin repeat protein also implicated in WNT pathway specification. Knockout of diversin in mice resulted in defective WNT-PCP signaling but hyper-activated signaling through the canonical WNT pathway (Allache et al., 2015; Jones et al., 2014) – phenotypes that are opposite to *usp9x* knockout but suggest a similar mode of pathway coupling. Importantly, diversin has also been reported to be overexpressed in breast cancer cells and contributes to both proliferation and invasion (Yu et al., 2014). Shared use of certain signaling factors – like dishevelled and diversin – between canonical and non-canonical WNT pathways may facilitate mechanisms of pathway specification based on inverse coupling – effectively operating as a “zero sum game”. For example, if a shared component like dishevelled or diversin is limiting for pathway activation then engagement by one pathway may consequently prevent activation of the coupled pathway. Thus, regulatory mechanisms that impact stability of shared factors or their interaction networks have strong potential to contribute to pathway specification.

DVL2 has been reported to undergo both positive and negative regulation by the ubiquitin system through the action of multiple different E3 ubiquitin ligases and deubiquitylating enzymes. For example, both ITCH and NEDD4L have been shown to negatively regulate DVL2 stability by conjugating K48-linked polyubiquitin chains on DVL2, targeting it to the proteasome for degradation (Ding et al., 2013; Liu et al., 2014; Marikawa and Elinson, 1998; Mukai et al., 2010; Tauriello et al., 2010; Wei et al., 2012). On the other hand, K63-linked polyubiquitin modifications of the DVL2 DIX domain were reported to enhance canonical WNT activation (Tauriello et al., 2010) while other reports have suggested these modifications may antagonize canonical WNT activation (Madrzak et al., 2015). The apparent regulation of DVL2 by multiple members of the NEDD4 family could result from functional redundancy, but could also reflect differential regulation based on expression in different contexts or different cell types. Our finding that siRNA targeting of NEDD4L or WWP1 partially rescued WNT activation in the absence of USP9X (**FIG 3E**) is consistent with a redundancy model, but it does not exclude the possibility that USP9X antagonizes multiple exclusive ubiquitylation events on DVL2 mediated by different E3 ubiquitin ligases. Indeed, our data indicates that functional regulation of DVL2 by USP9X involves reversal of multiple distinct ubiquitylation events (**FIG 4B**) – possibly mediated by multiple E3 ubiquitin ligases – since no single K→R mutation was capable of suppressing the loss of WNT activation phenotype observed in the absence of USP9X (**FIG S4F**). Furthermore, the *in vitro* analysis of USP9X activity presented here suggests that USP9X may be unique in its ability to operate on WWP1-ubiquitylated DVL2, and that it may have activity toward atypical ubiquitin polymers (**FIG 2B-D**). Additional biochemical and genetic studies will be required to fully elucidate how USP9X affects various distinct ubiquitylation events on DVL2 that may be mediated by different E3 ubiquitin ligases.

Previous studies have reported conflicting roles for USP9X with respect to canonical WNT activation. Specifically, USP9X has previously been reported to positively regulate the stability of β-catenin in glioma cells (Yang et al., 2016). However, USP9X was also reported to negatively regulate the stability of β-catenin in mouse neural progenitor cells (Premarathne et al., 2017). Our studies are consistent with a positive role for USP9X in canonical WNT pathway activation and our data indicate that USP9X functions upstream of the destruction complex (**FIG S4A**). Although our findings are consistent with a role for USP9X in the regulation of WNT at the level of DVL2 we cannot exclude the possibility of a context-dependent role for USP9X at other tiers of the WNT pathway. Furthermore, it is possible that expression of different NEDD4 family members, other USP9X-interactors, or substrate adaptors and targeting factors could modify USP9X function in a context-specific manner. Although previous studies have reported that Wnt5a-mediated activation of the WNT-PCP pathway in human breast cancer cells causes increased cell migration as a result of DVL2/Daam1-dependent activation of RhoA (Luga et al., 2012; Zhu et al., 2012) USP9X has not previously been linked to the regulation of these activities. Data presented in this study indicates that USP9X and WWP1 interact to regulate the ubiquitylation status of DVL2, which impacts DVL2 localization to actin-rich projections, Rho activation, and motility in breast cancer cells. These findings reveal a novel mode for regulation of WNT-PCP activation and motility of human breast cancer cells.

### USP9X: a Complex Factor in Cancer

Given that USP9X has been implicated in the regulation of multiple receptor signaling and trafficking pathways, it is not surprising that in recent years USP9X dysregulation has been linked to many different types of cancers. USP9X has been suggested to exhibit oncogenic behavior in some cancer contexts including multiple myeloma, melanoma, lung cancer, and breast cancer (Kushwaha et al., 2015; Li et al., 2017; Potu et al., 2017; Schwickart et al., 2010). In contrast, USP9X has also been proposed to have tumor suppressor activity in other cancers including pancreatic ductal adenocarcinoma (Pérez-Mancera et al., 2012) and colorectal adenocarcinoma (Khan et al., 2018). Recently, USP9X has been shown to be required for the attenuation of EGFR signaling (Savio et al., 2016) which is suggestive of a tumor suppressor function. Taken together, these reports suggest a complex role for USP9X in cancer progression that may be highly context-dependent. Our findings suggest that USP9X functions together with WWP1 to regulate a ubiquitin rheostat on DVL2. The ubiquitylation status of DVL2 determines its engagement with either canonical or non-canonical WNT factors. Based on these findings we hypothesize that in some cellular contexts increased USP9X activity may sensitize cells to activation of the canonical WNT pathway (**FIG 3D**), which promotes proliferation and tumor growth, while simultaneously antagonizing the non-canonical WNT-PCP pathway which is involved in cell motility and metastasis (**FIG 6C-F**). These results indicate that dysregulation of USP9X can function to promote canonical WNT-driven proliferation while simultaneously guarding against migration and metastasis mediated by the WNT-PCP pathway. Based on these findings, we propose that small molecule inhibitors of USP9X may be effective as antagonists of WNT-based proliferation (**FIG 3C**) but they may also ultimately promote migration and metastasis (**FIG S6B**).

### Regulation of cell signaling pathways by DUB-E3 interactions

The ubiquitin proteasome system contributes to the tight spatial and temporal regulation of protein trafficking, localization, and stability. Recently, several examples of interactions between E3 ubiquitin ligases and DUBs have been reported, raising questions about the regulatory significance and design logic of such interactions. In principle, DUB-E3 interactions could facilitate a number of useful design features, including E3 antagonization, E3 deubiquitylation, or polyubiquitin chain editing on substrates. E3 antagonization has the potential to result in a futile cycle, but coordinated regulation of ligase and DUB activities could also establish a ubiquitin rheostat on a shared substrate. Such a rheostat has been reported to occur in the context of ER quality control machinery, where DUB activities can sharpen the distinction between folded and misfolded membrane proteins (Zhang et al., 2013). Another striking example involves the DUB USP7, which was shown to interact with the MAGE-L2-TRIM27 E3 ubiquitin ligase to regulate the ubiquitylation status of WASH, a shared substrate, on the endosome (Wu et al., 2014). Interestingly, USP7 has been reported to interact with multiple E3 ubiquitin ligases (Kim and Sixma, 2017) but the functional significance of many of these interactions remain unclear. E3-DUB interactions have also been reported to protect E3 ubiquitin ligases from degradation, thus promoting E3 activity. For example, the DUB USP19 promotes cellular proliferation by stabilizing KPC1 (also known as RNF123), a ubiquitin ligase for the cyclin-dependent kinase inhibitor p27Kip1 (also known as CDKN1B) (Lu et al., 2009). A striking example of polyubiquitin chain editing occurs with the protein A20, a potent inhibitor of the NFkB signaling pathway that contains both a DUB domain and an E3 ubiquitin ligase domain (Wertz et al., 2004). The DUB domain of A20 removes a K63-linked polyubiquitin chain from the Receptor Interacting Protein (RIP), a protein essential for NFkB signaling. Subsequently, the E3 ubiquitin ligase domain of A20 conjugates a K48-linked polyubiquitin chain to RIP targeting it for degradation by the proteasome – thus inhibiting the NFkB pathway. Thus, there are numerous examples of E3-DUB interactions that establish ubiquitin rheostats on substrates, protect and stabilize E3 ubiquitin ligases, and coordinate activities to achieve polyubiquitin chain editing in signaling complexes.

We have identified a novel E3-DUB interaction that functions to provide layered regulation of the WNT pathway. Our results indicate that the activities of WWP1 and USP9X are antagonistic and function to establish a ubiquitin rheostat on DVL2 that is a critical regulator of WNT pathway specification. Functionally, USP9X antagonizes WWP1 activity towards DVL2 (**FIG 2C-D**), although our data cannot exclude the possibility that WWP1-USP9x interactions function in a chain-editing capacity on polyubiquitinated DVL2. Ultimately, rigorous biochemical analysis dissecting mechanisms that regulate the balance of different ubiquitylation and deubiquitylation activities toward DVL2 will be critical to understand how its specification for either canonical or noncanonical WNT signaling pathways is determined.

## STAR Methods section

### CONTACT FOR REAGENT AND RESOURCE SHARING

Further information and requests for resources and reagents should be directed to and will be fulfilled by the Lead Contact, Jason MacGurn.

### EXPERIMENTAL MODEL AND SUBJECT DETAILS

#### Cell lines

1. **<u>HEK239 cells</u>** Source: American Type Culture Collection (ATCC), Catalog #: CRL-1573 Culture/growth conditions: Cultured in DMEM with 10% FBS and 1% penicillin/streptomycin at 37°C in 5% CO_2_ Sex: Female Authentication: Yes (ATCC authenticated)
2. **<u>MDA-MB-231 cells</u>** Source: ATCC, Catalog #: HTB-26 Culture/growth conditions: Cultured in RPMI with 10% FBS and 1% penicillin/streptomycin at 37°C in 5% CO_2_ Sex: Female Authentication: Yes (ATCC authenticated)
3. **<u>HEK293 STF cells</u>** Source: ATCC, Catalog #: HEK 293 STF (ATCC® CRL-3249™) Culture/growth conditions: Cultured in RPMI with 10% FBS and 1% penicillin/streptomycin at 37°C in 5% CO_2_ Sex: Female Authentication: Yes (ATCC authenticated)

### METHOD DETAILS

#### Assays for measuring WNT activation

Topflash reporter assays in HEK293 STF cells were performed as follows: HEK293 STF cells were seeded on 24-well plates at ∼50% confluency. Cells were treated with 100ng/mL purified recombinant WNT3a (R&D Systems) and 100ng/mL purified recombinant R-spondin (R&D systems) 24 hours prior to lysis with 1X Passive Lysis buffer (Promega). Luciferase activity was measured using the Steady-Glo luciferase Assay (Promega). Luciferase signal was normalized to viable cell number using Cell-Titer Glo Assay (Promega). For siRNA knockdown studies, cells were treated for 72 hours prior to lysis, for inhibitor studies cells were treated for 24 hours prior to lysis, and for overexpression studies cells were transfected with plasmid DNA for 48 hours prior to lysis.

Topflash reporter assays in MDA-MB-231 cells were performed as follows: Cells were plated, co-transfected with TOPflash and Renilla expression plasmids at 24 hours and lysed at 48 hours using the Dual-Glo Luciferase Assay system (Promega). Luciferase signal was normalized to co-transfected Renilla expression. Data were normalized to a positive control (set at 100% activation). Assays were performed in triplicate and repeated at least 3 times.

Β-catenin stabilization assay: HEK293 STF cells were cultured in Dulbecco’s modified Eagle’s Medium (DMEM) and activated with WNT3A ligand or LiCl for 24 hours. Then cells were lysed and nuclear/cytoplasmic fractionation was performed as previously described (Thorne et al., 2010).

#### Transfections

Plasmid and siRNA transfections were performed using Lipojet (SignaGen, SL100468) and Lipofectamine LTX with PLUS reagent (Thermo Fisher, 15338100) according to manufacturer’s protocol.

#### Immunoblots

Whole-cell lysates were generated by lysing cells in Laemmli Sample Buffer (Bio-Rad) with 5% βME and subsequently boiling samples at 95°C for 5 minutes. Proteins were analyzed by SDS-PAGE and immunoblotting using the LiCor Odyssey reagents and protocols. Fluorescence signaling was detected using an Odyssey CLx Infrared Scanner (LiCor).

#### Cell lines

HEK293 STF cells and MDA-MB-231 cells were purchased from the American Type Culture Collection (ATCC). HEK293 STF cells were cultured in DMEM with 10% FBS and 1% penicillin/streptomycin at 37°C in 5% CO2. MDA-MB-231 cells were cultured in DMEM with 10% FBS and 1% penicillin/streptomycin at 37°C in 5% CO2. MDA-MB-231 cells stably expressing FLAG-WWP1, FLAG-DVL2, FLAG-DUB-DVL2, or FLAG-DUB*-DVL2 were generated using the pQCXIP retroviral vector system. MDA-MB-231 *usp9x* knockout cells were generated using a CRISPR/Cas-9 genome editing system.

#### SILAC-based Quantitative Proteomic Analysis

Quantitative mass spectrometry analysis by SILAC was performed on MDA-MB-231 cells stably expressing the indicated FLAG-tagged substrates. These cells were cultured in the presence of either heavy or light isotopes (lysine and arginine) for at least 6 doublings to ensure incorporation of these isotopes into 100% of the proteome. Affinity purification was performed as previously described (Lee et al., 2017). Eluted proteins were digested with 1μg Trypsin Gold (Promega) at 37 F overnight. Digested peptides were cleaned-up on a Sep-Pak C18 column. Purified peptides were then dried, reconstituted in 0.1% trifluoroacetic acid, and analyzed by LC-MS/MS using an Orbitrap XL mass spectrometer. Database search and SILAC quantitation were performed using MaxQuant software.

#### Co-Immunoprecipitation Studies

For coimmunoprecipitation assays, cells were washed with cold PBS and lysed with a nondenaturing lysis buffer (NDLB): 50mM tris (pH 7.4), 300mM NaCl, 5mM EDTA, 1% Triton X-100, cOmplete protease inhibitor cocktail (Roche), Phosphatase inhibitor cocktail (Roche), 1mM PMSF, 1mM Phenanthroline, 10mM Iodoacetamide, and 20μM MG132. Lysates were diluted to 1mg/mL with NDLB and incubated with Sigma FLAG EZ-view beads for 2 hours with rotation at 4°C. Beads were washed 3 times with cold NDLB and once with cold PBS. Proteins were then eluted from beads using sample buffer and samples were processed for SDS-PAGE followed by immunoblotting.

#### Fluorescence Microscopy

MDA-MB-231 cells were seeded on coverslips at 50% confluency. At 24 hours after seeding, cells were transfected with FLAG-Dvl2 using LipoJet Transfection Reagent (Signagen). 72 hours after seeding cells were fixed and permeabilized using 4% paraformaldehyde (PFA) and immunofluorescence block buffer (PBS + 0.1% Triton-x + 10% FBS). Cells were then labeled with primary antibodies for 1 hour followed by a 1 hour incubation with alexa-fluor conjugated secondary antibodies. Coverslips were mounted on slides using ProLong AntiFade mountant (Thermo). Microscopy images were acquired using a DeltaVision Elite system (GE Healthcare) and processed using SoftWoRx software (GE Healthcare, Chicago, IL).

#### In Vitro Ubiquitin Conjugation and Deubiquitylation Assays

Due to the limited solubility of full-length DVL2 in solution, in all *in vitro* conjugation and deconjugation assays HA-DVL2 was purified from human HEK293 cells using magnetic anti-HA beads (Pierce #88836) and all conjugation/deconjugation reactions were performed on HA-DVL2 bound to anti-HA beads. Lysis and purification were performed as follows: 24 hours after seeding, HEK293 cells were transfected with HA-Dvl2 and 48 hours post-transfection cells were lysed in NDLB and clarified by centrifugation. Lysates bound anti-HA magnetic beads overnight at 4°C with rotation. Beads were washed 3 times in NDLB and 1 time in cold 1X PBS before being resuspended in conjugation buffer (40 mM Tris pH 7.5, 10 mM MgCl2, and 0.6 mM DTT) for immediate use in conjugation/deconjugation reactions.

For the conjugation reactions, 56 nM GST-Ube1 (Boston Biochem, Cambridge, MA, E-300), 0.77 mM UBE2D3, 2.3 mM Ub, HA-DVL2, and 60 nM FLAG-WWP1 (Sigma, SRP0229) were incubated in conjugation buffer (40 mM Tris pH 7.5, 10 mM MgCl2, and 0.6 mM DTT) with 1 mM ATP, at 37°C. Reactions were initiated by the addition of FLAG-WWP1. Samples were removed at indicated time-points, boiled in 2x Laemmli sample buffer for 10 min, and analyzed by blotting for HA-DVL2 with DVL2 antibody (Cell Signaling Technology, #3216) using goat anti-rabbit secondary (IRDye 800 CW, LI-COR Biosciences, Lincoln, NE). Quantification was performed using ImageStudioLite software (LI-COR).

For deconjugation reactions, polyubiquitinated substrate (HA-DVL2) was generated using the conjugation assay described above within one exception in that 0.1 mM ATP was used. The conjugation reaction was stopped by incubation with 0.75 units Apyrase in a 90 µl reaction at 30°C for 1 hr. For the deconjugation reaction, 100 nM His6-USP9X (Boston Biochem E-552), 100nM OTUB1, or 100nM AMSH was added to conjugated HA-DVL2 as well as 10X DUB buffer (0.5 M Tris pH 7.5, 500 mM NaCl, 50mM DTT) to 1X. Reactions were initiated by the addition of His6-USP9X protein. The reaction was incubated at 37°C. Samples were removed at indicated time-points and boiled in 2x Laemmli sample buffer. HA-DVL2 levels were determined as described in the conjugation assay.

For Rheostat experiments, HA-DVL2 substrate was resuspended in conjugation buffer with 1mM ATP and treated with 56 nM GST-Ube1, 0.77 mM UBE2D3, 2.3 mM Ub, and indicated concentrations of FLAG-WWP1 and His-USP9X for 60 minutes at 37°C. Samples were then boiled in 2x Lammli sample buffer for 10 minutes and analyzed by blotting for HA-DVL2 with DVL2 antibody (CST #3216). Line profile analysis was performed using ImageJ (NIH).

#### Migration Assays

Cell culture dishes were coated with 5 μg/ml rat tail ColI (Corning #354236) in PBS for 1 h at 37°C then cells were plated at low density and incubated at 37°C for 1 hour in cell culture medium. Cells were kept at 37°C in SFM4MAb medium (HyClone, Logan, UT) supplemented with 2% FBS at pH 7.4 during imaging using phase-contrast and fluorescence microscopy techniques. Images were acquired at 5 min intervals for 6 h, using DeltaVision Elite system software (GE Healthcare) and cell speed was quantified using SoftWoRx software (GE Healthcare, Chicago, IL). Wind rose plots were generated by transposing x,y coordinates of cell tracks to a common origin.

#### Rho Activation Assays

Rho activation assays were performed using the Rho Activation Assay Kit (Millipore Sigma #17-924). MDA-MB-231 cells (WT and *usp9x* KO cells) were grown until confluent, washed in ice-cold 1XPBS, lysed in ice-cold 1X Mg^2+^ Lysis/Wash buffer (MLB) (Millipore Sigma #20-168), scraped and collected in microfuge tubes on ice, and incubated at 4°C for 15 minutes with agitation. Lysates were then cleared of cellular debris by centrifugation at 14,000xg for 5 minutes at 4°C. Clarified lysate was then bound to Rho Assay Reagent (Millipore Sigma #14-383) at 4°C for 45 minutes with gentle agitation. Agarose beads were then washed 3 times with 1X MLB and proteins were eluted in 2x Laemmli reducing sample buffer (BioRad #1610737) by boiling at 95°C for 5 minutes. Samples were fractionated by SDS-PAGE and analyzed by blotting using the Anti-Rho (-A, -B, -C) clone 55 antibody (Millipore Sigma #05-778).

GTPγS/GDP loading for positive and negative controls were prepared as described above with an additional 30 minute incubation at 30°C with agitation of cell lysate in the presence of 10mM EDTA and either 100uM GTPγS (Millipore Sigma #20-176) or 1mM GDP (Millipore Sigma #20-177). This step was performed prior to binding the Rho Assay Reagent.

### QUANTIFICATION AND STATISTICAL ANALYSIS

#### Statistical Analysis

Detailed statistical analysis can be found in the accompanying Statistical Reporting Documents for each figure. For each statistical analysis a Student’s T-test was used to test for a statistically significant difference between the means of the two variables of interest. The alpha value for each experiment was set at 0.05 and a p-value was calculated using the Student’s t-test function in Microsoft Excel to determine statistical significance. Each figure with statistical analysis represents n≥3 for all coIP and luciferase assay experiments, where n represents biological replicates. For quantitation of all microscopy-based data an n≥25 was used where n represents single cells. In all figures including statistics, error bars represent standard deviation of the mean. Details of statistical analysis can also be found in figure legends.

### KEY RESOURCES TABLE

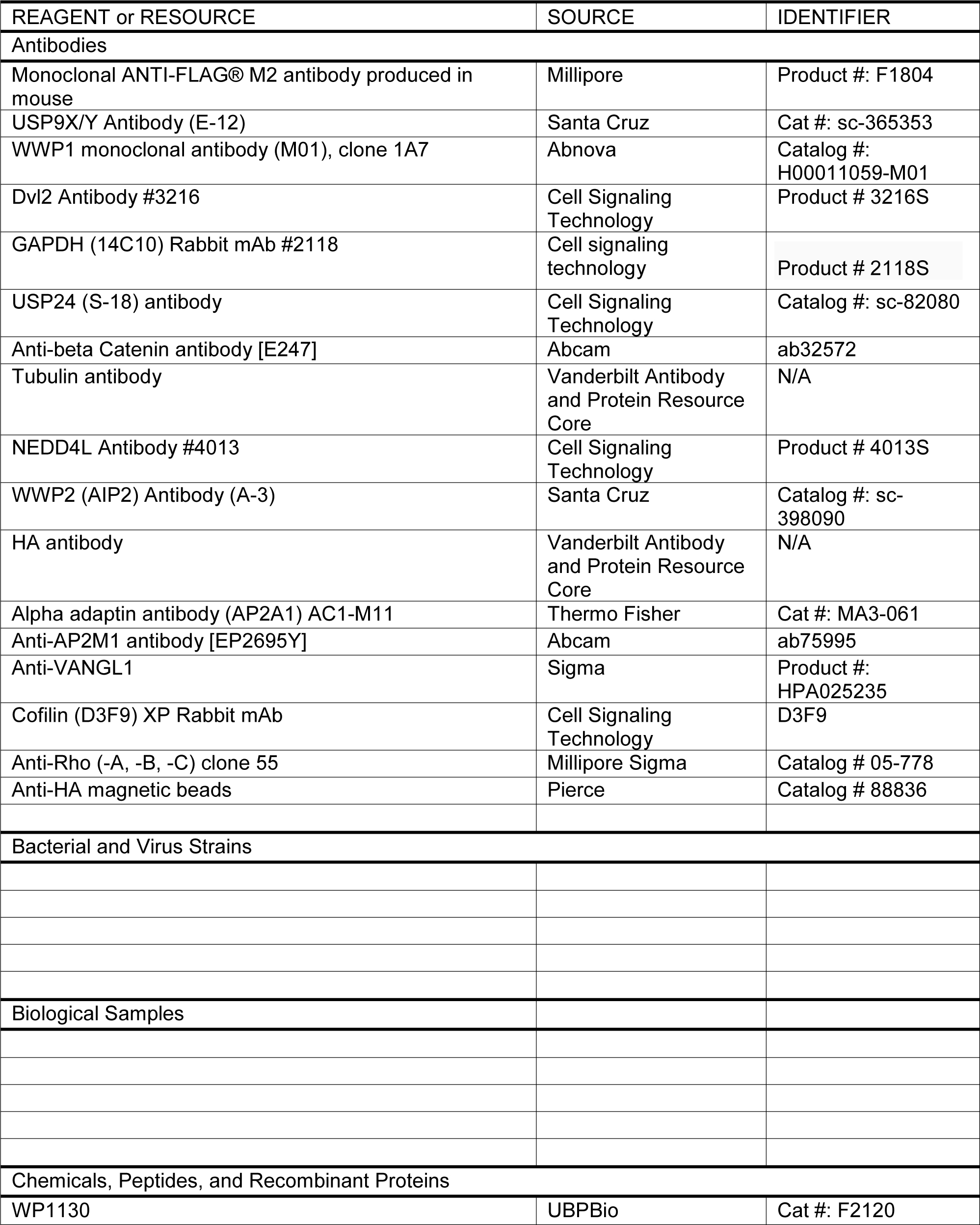
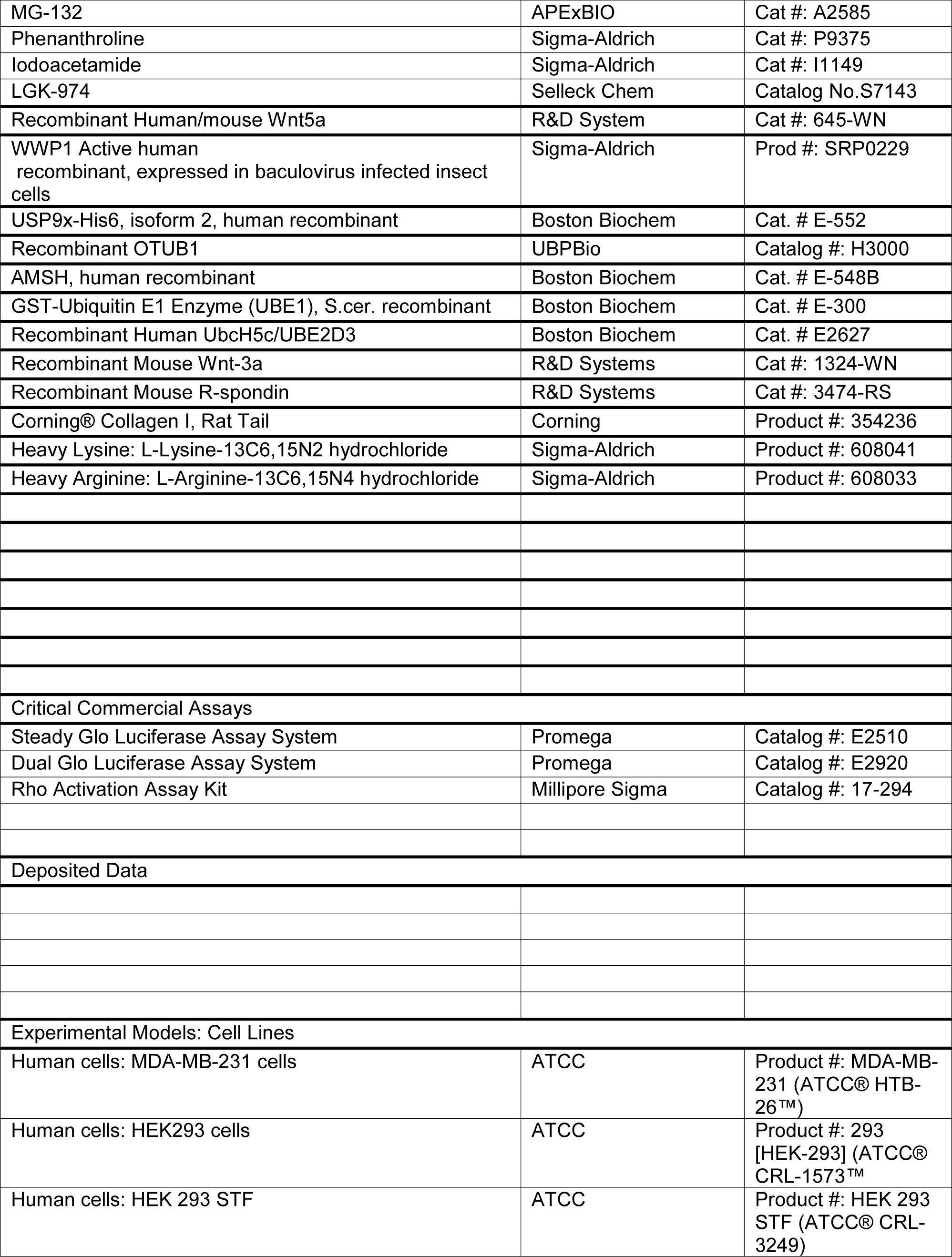
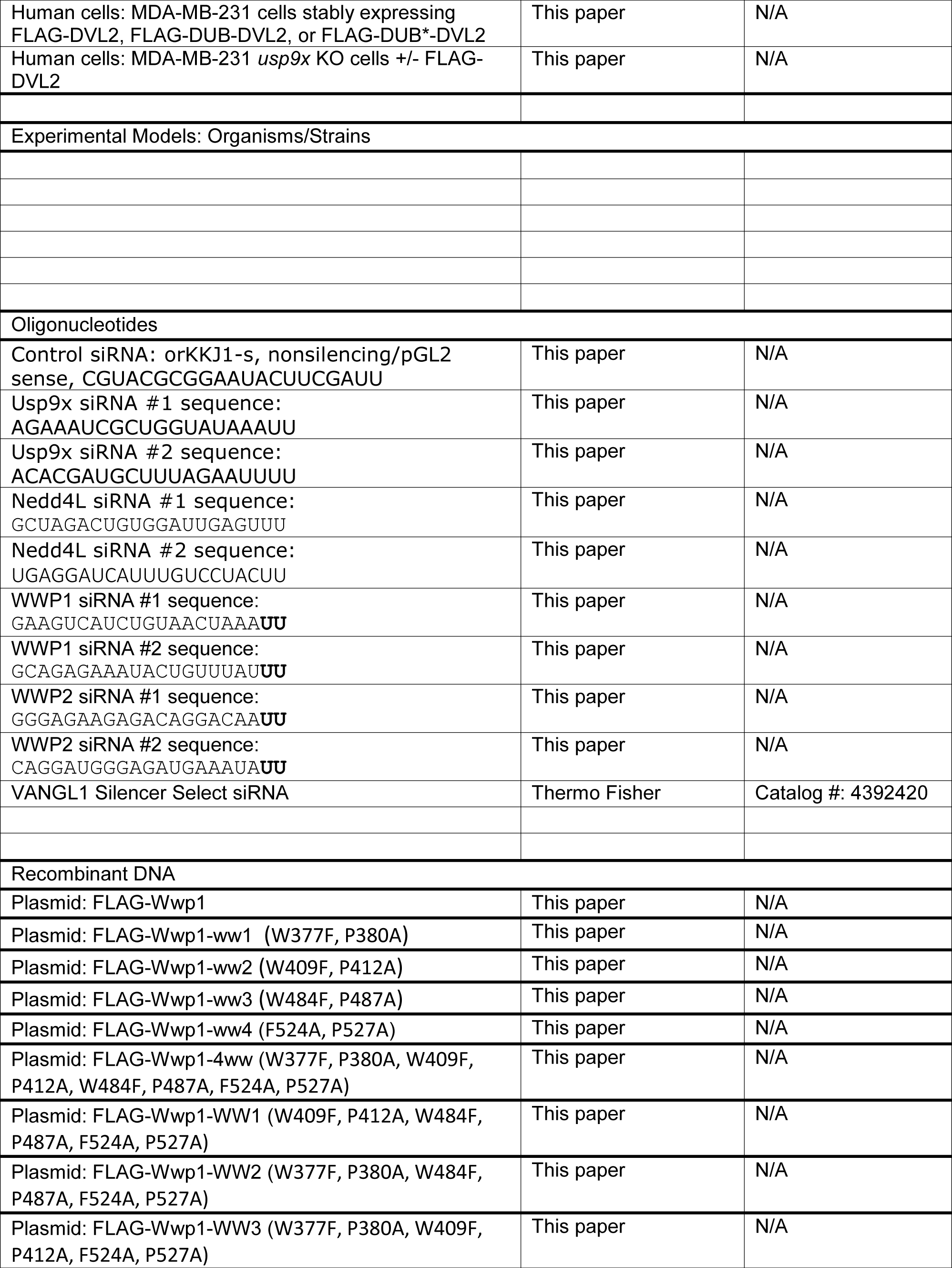
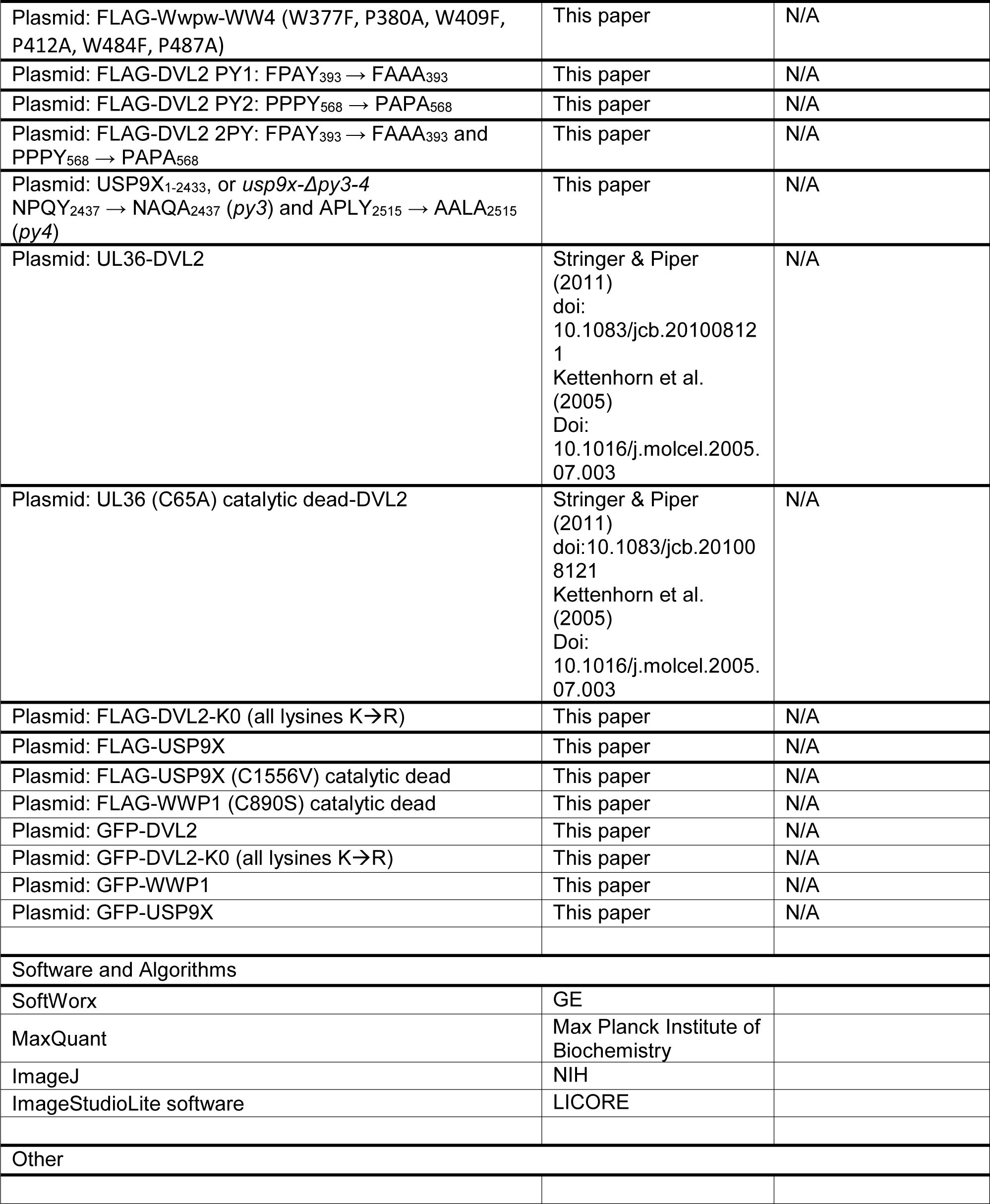

## Acknowledgements

We are very grateful to K. Rose for helpful advice regarding technical aspects and analysis of quantitative proteomic data. We are also grateful to E. Lee, K. Saito-Diaz, V. Ng for reagents and technical advice. We also thank T. Graham, S. M. Ahmed, S. Lee, and N. Hepowit for critical reading of this manuscript. This research was supported by NIH grant R00 GM101077 (to J.A.M.) and GI SPORE grant in GI Cancer (P50 CA095103).

---

**Supplemental Figure S1.**
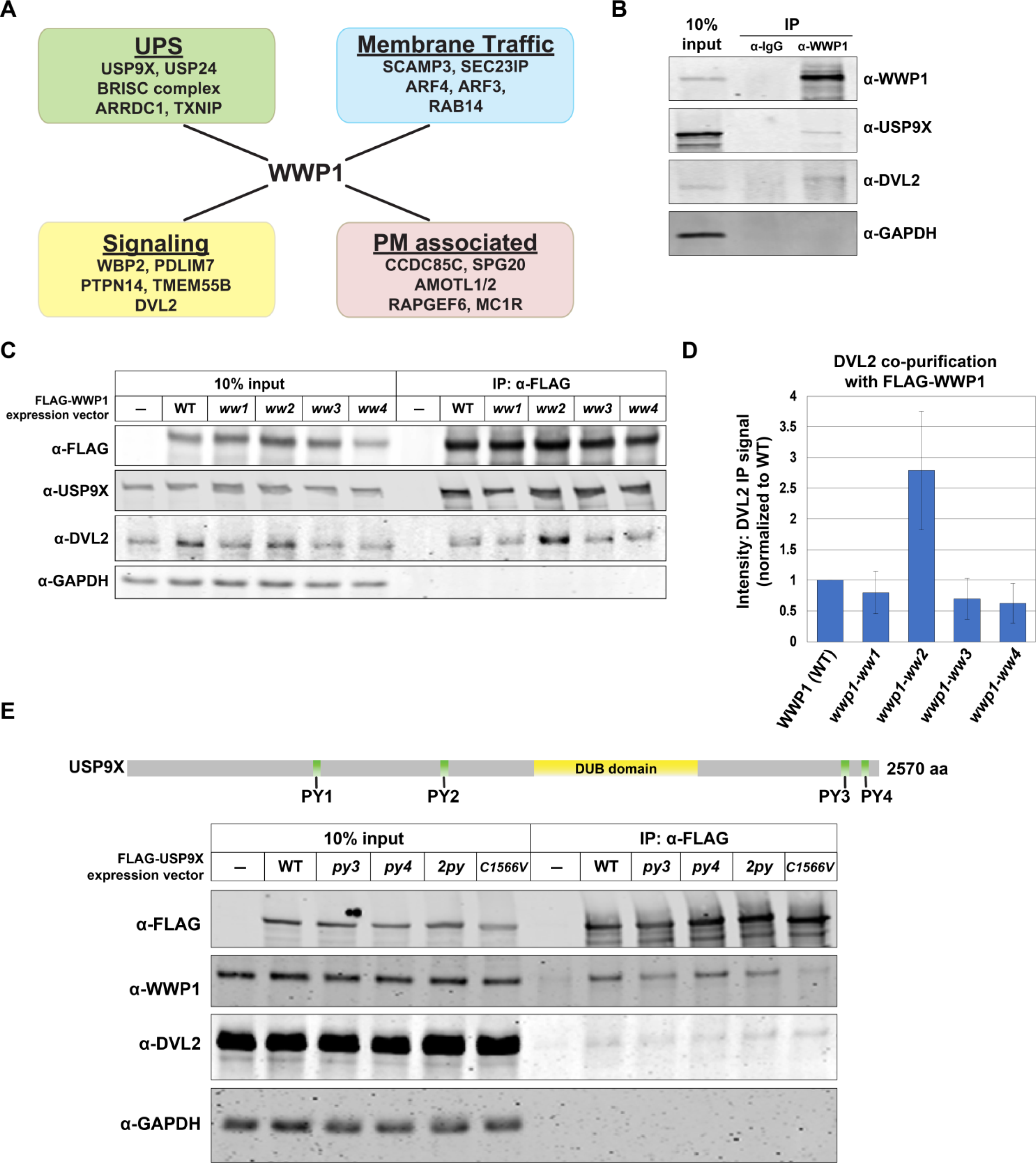
Related to Figure 1. USP9X, WWP1, and DVL2 interactions are scaffolded by WW-PY interactions **(A)** Schematic summarizing putative WWP1 interacting proteins identified from SILAC-MS. Complete data set for this experiment is reported in Table S1. **(B)** Endogenous WWP1 was immunoprecipitated from MDA-MB-231 cells. Input (10%) and immunoprecipitates were resolved by SDS-PAGE and immunoblotted for the indicated species. **(C)** Analysis of co-purification of DVL2 and USP9X with WWP1 variants. The indicated WWP1 variants were FLAG affinity purified, and quantitative immunoblots were performed to assess co-purification of interacting factors. Mutant variants of WWP1 included point mutations disrupting the ability of individual WW domains to bind PY motifs. **(D)** Quantification of DVL2 co-purification with FLAG-WWP1 variants (normalized to WT) over multiple experiments (n=3). **(E)** Analysis of co-purification of WWP1 and DVL2 with USP9X mutant variants. The indicated mutant USP9X variants were FLAG affinity purified, and quantitative immunoblots were performed to assess co-purification of interacting factors. *py3* and *py4* mutations indicate point mutations in PY3 and PY4, respectively, while *2py* indicates both C-terminal PY motifs were mutated. *C1566V* is mutated at the catalytic cysteine residue of USP9X.

**Supplemental Figure S2.**
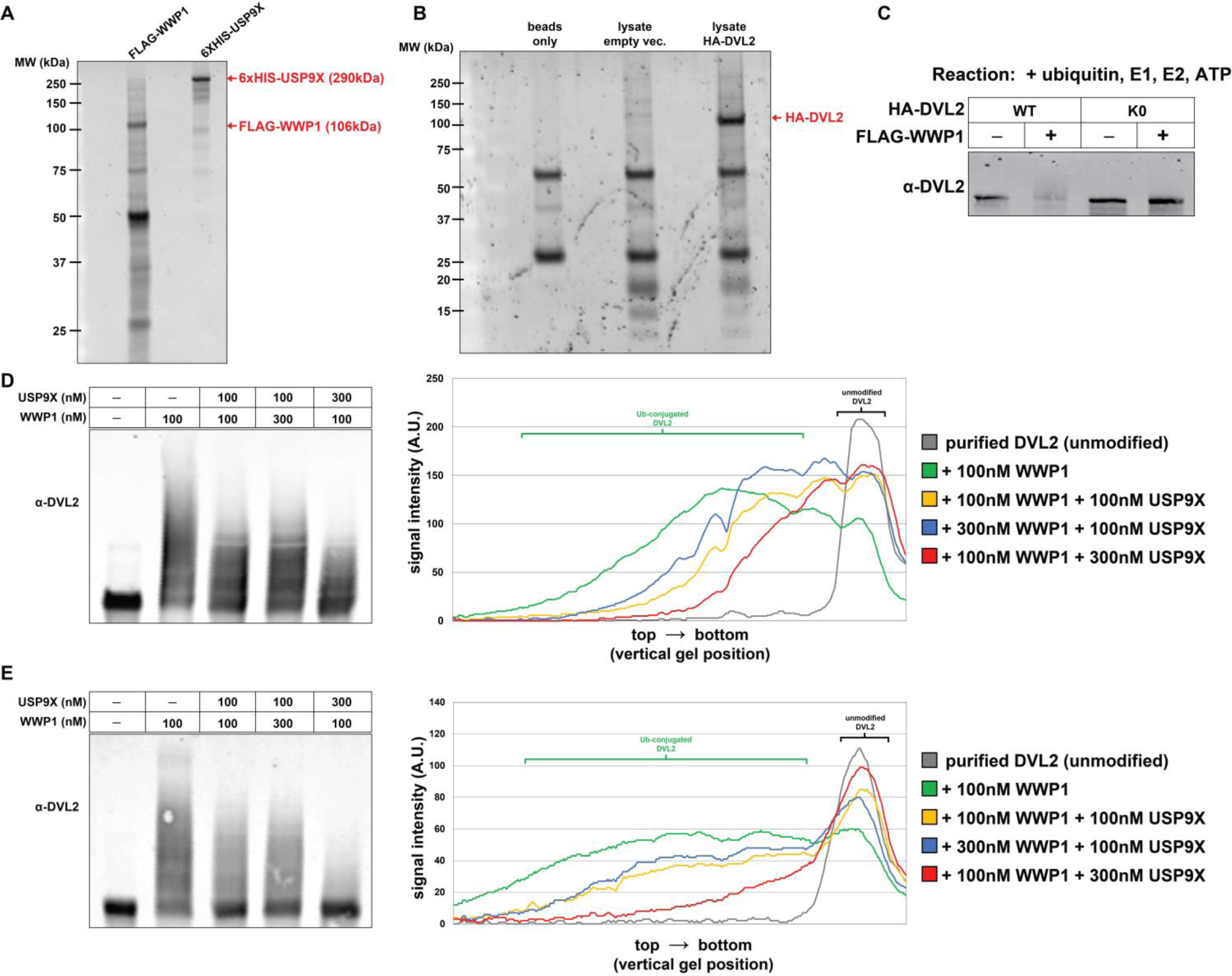
Related to Figure 2. Analysis of WWP1 and USP9x activity toward DVL2 *in vitro*. **(A)** Purity of recombinant FLAG-WWP1 and 6XHIS-USP9X used in *in vitro* conjugation/deconjugation reactions (FIG 2) shown by SYPRO Ruby gel staining. **(B)** Purity of HA-DVL2 (purified from cultured cells) used in *in vitro* conjugation/deconjugation reactions (FIG 2) shown by SYPRO Ruby gel staining. **(C)** *In vitro* ubiquitylation reactions were performed using recombinant purified FLAG-WWP1 (60nM) and HA-DVL2 (wildtype and K0, a variant where all lysine residues are mutated to arginine) purified from cultured cells. E3 conjugation reactions were allowed to proceed for 60 minutes before the conjugation reaction was terminated. **(D-E, left panels)** Biological replicates of *in vitro* ubiquitylation rheostat reaction shown in FIG 2C-D, experiments were performed using indicated combinations of purified FLAG-WWP1 and 6XHIS-USP9X with HA-DVL2 purified from cultured cells. Reactions were allowed to proceed for 60 minutes before reaction was terminated and HA-DVL2 was resolved by SDS-PAGE and immunoblot. (**D-E, right panels**) Quantification of HA-DVL2 ubiquitylation under indicated conditions using ImageJ line profile analysis of immunoblot.

**Supplemental Figure S3.**
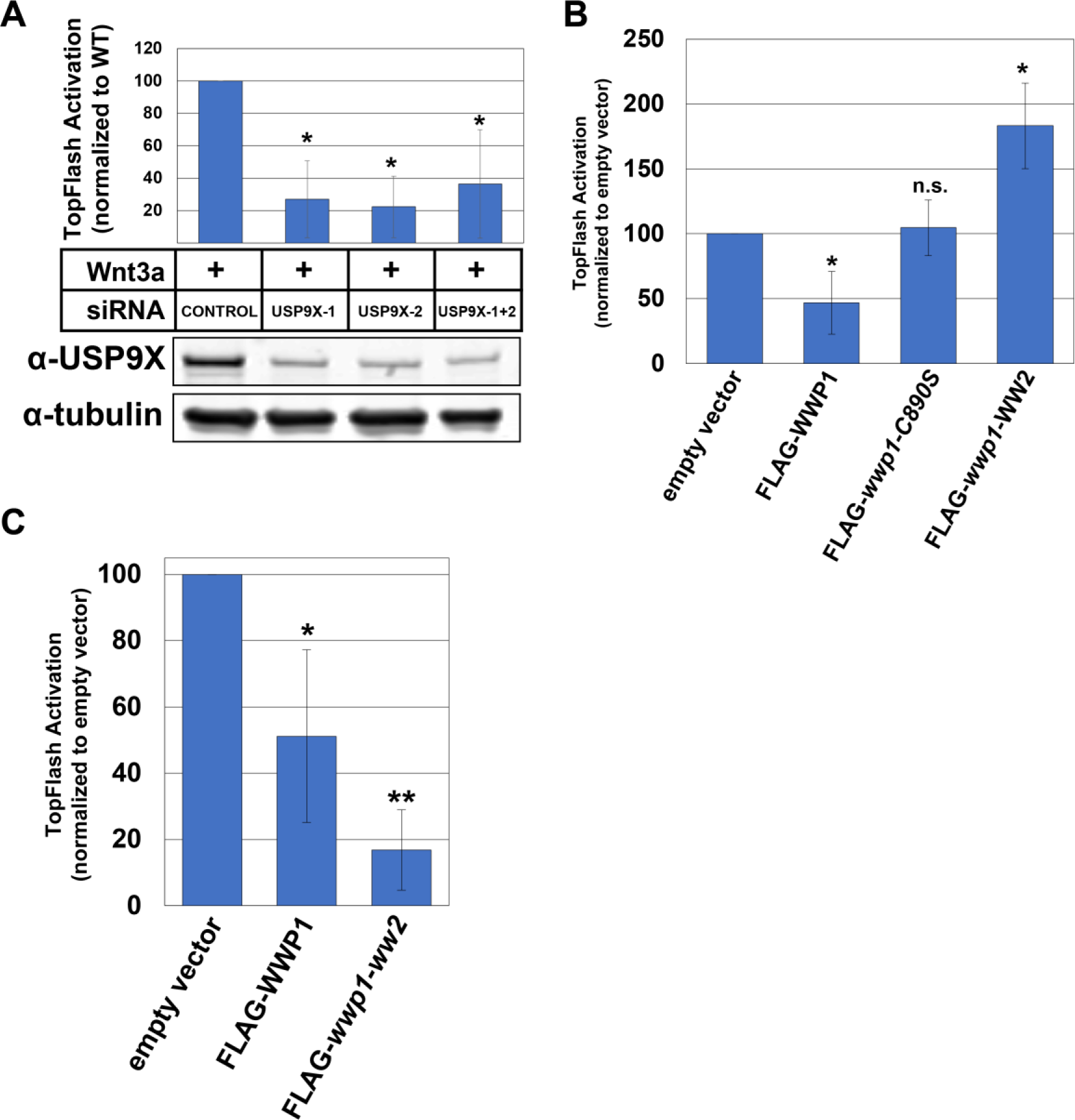
Related to Figure 3. USP9X and WWP1 regulate canonical WNT activation. **(A)** Analysis of ligand-stimulated WNT activation in the presence of control siRNA or siRNAs targeting USP9X knockdown. TopFLASH luciferase assays were used to measure WNT activation (top panel) and immunoblotting was performed to assess knockdown efficiency (bottom panel). The asterisk indicates significant difference compared to control siRNA treatment (p<0.05). **(B)** TopFLASH activation of STF-293 cells transfected with empty vector or plasmids for CMV-driven expression of wildtype (WT) FLAG-WWP1, *wwp1-C890S* (a catalytic dead variant) or *wwp1*-WW2 (a variant with only WW2 intact) (see **FIG 1E** and **1F**). The asterisk indicates a statistically significant difference compared to the empty vector control (p<0.05) while n.s. indicates no significant difference compared to the empty vector control. **(C)** TopFLASH activation of STF-293 cells transfected with empty vector or plasmids for CMV-driven expression of wildtype (WT) FLAG-WWP1 or a *ww2* mutant variant (see **FIG S1C** and **S1D**). Single asterisk indicates statistically significant difference compared to empty vector control (p<0.05), while double asterisk indicates statistically significant difference compared to both empty vector control and wildtype WWP1 overexpression (p<0.005).

**Supplemental Figure S4.**
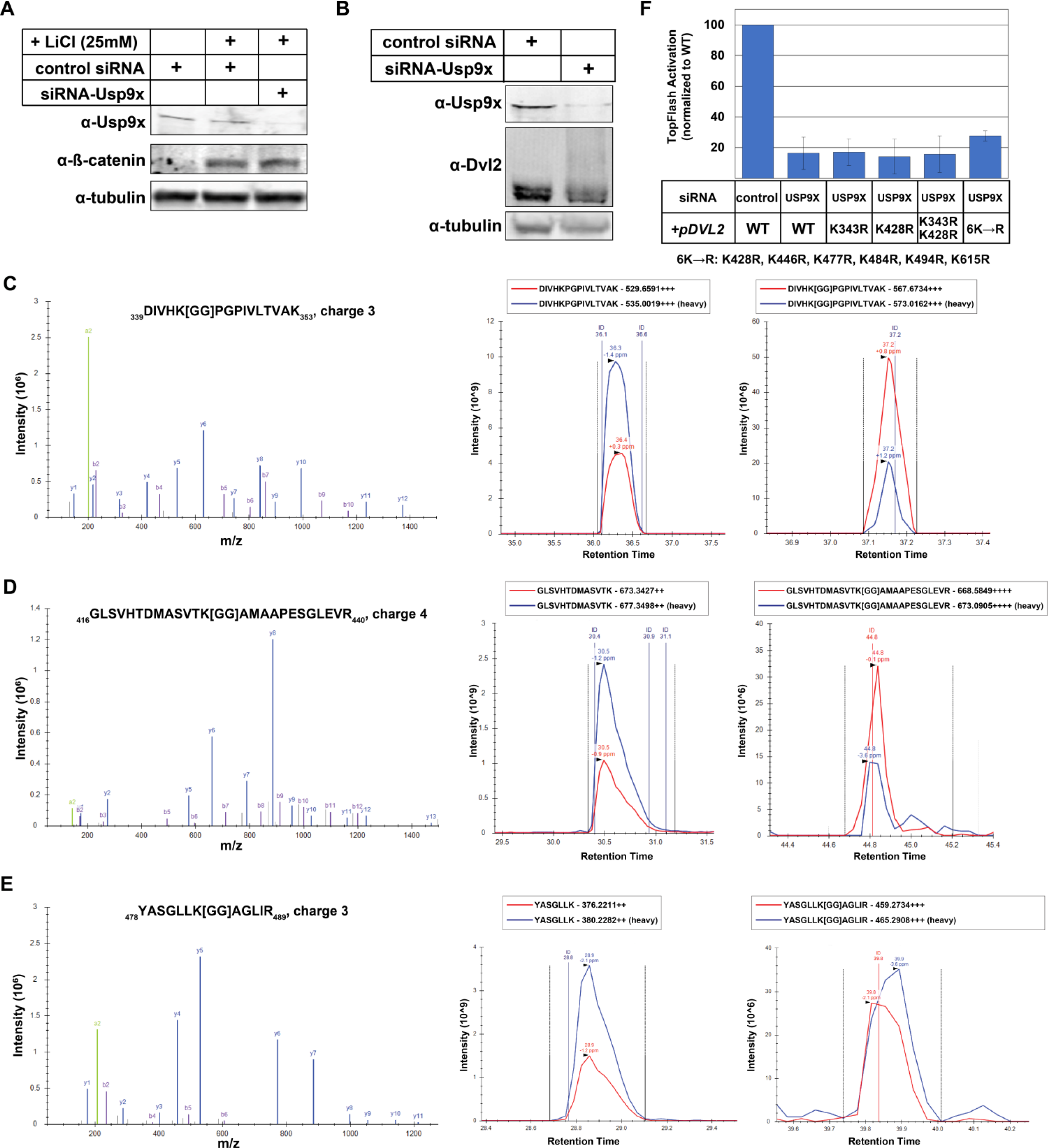
Related to Figure 4. USP9X regulates DVL2 ubiquitylation state. **(A)** Analysis of LiCl stimulation of WNT activation following transfection with control siRNA or siRNAs targeting USP9X knockdown. Immunoblotting was performed to assess knockdown efficiency and stabilization of nuclear β–catenin. **(B)** Immunoblotting analysis of DVL2 mobility by SDS-PAGE following transfection with control siRNA or siRNAs targeting USP9X knockdown. **(C-E)** Raw data for SILAC quantification (corresponding to **FIG 4B**) of individual ubiquitylation events on DVL2 including diGly modification of K343 **(C)**, K428 **(D)** and K484 **(E)**. MS2 spectra for each diGly peptide is shown in the left panel. To the right of each MS2 spectra, chromatograms are shown for heavy and light unmodified (middle) and diGly-modified (right) peptides, illustrating the quantification of each peptide. In each case, the diGly-modified form of the peptide is significantly elevated in the light sample, which is derived from the *usp9x* mutant sample (see **FIG 4B**). **(F)** Functional analysis of DLV2 lysine mutants. STF-293 reporter cells were transiently transfected with the indicated FLAG-DVL2 plasmids and siRNAs and analyzed for WNT activation using the TopFLASH luciferase assay.

**Supplemental Figure S5.**
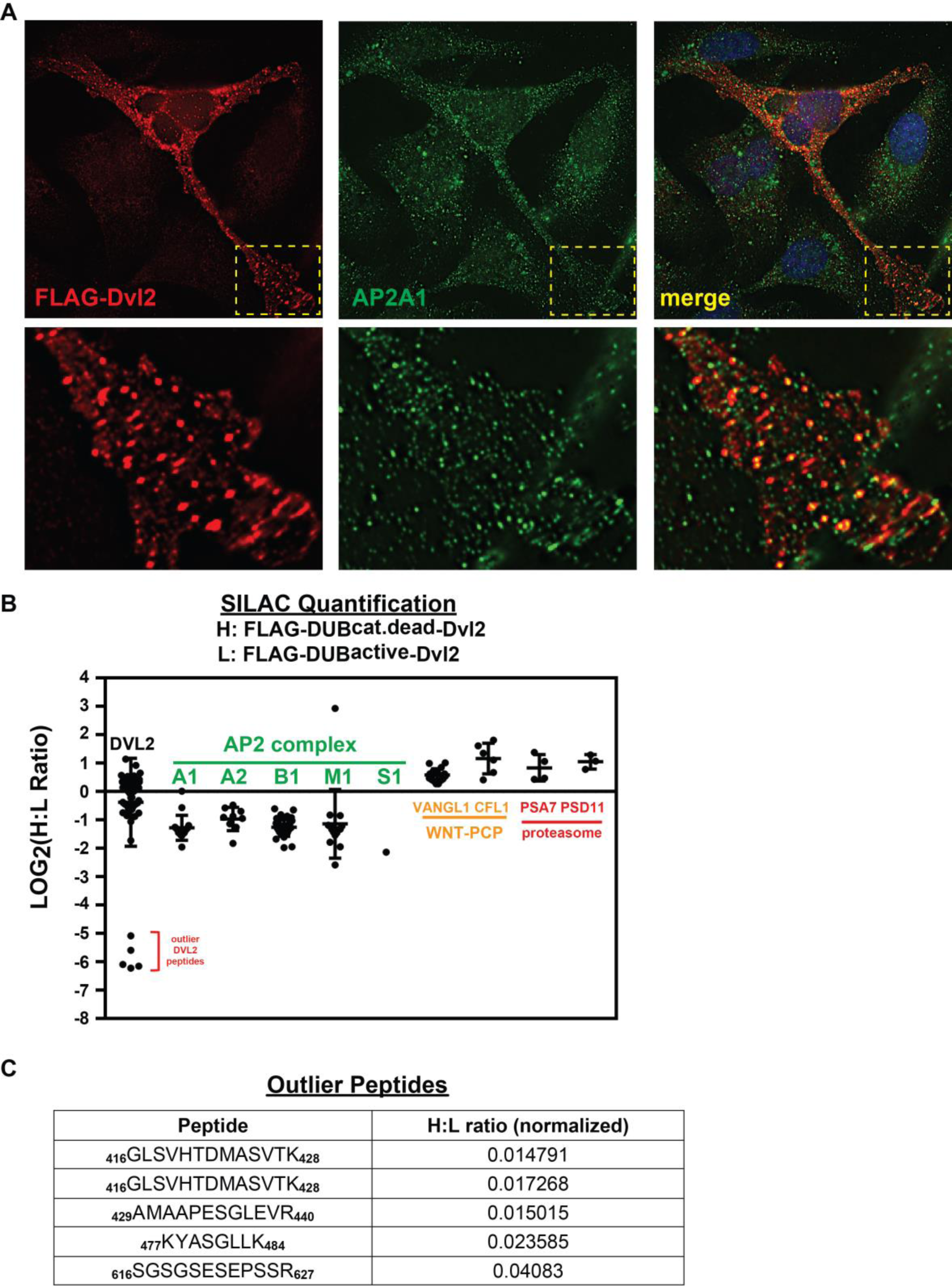
Related to Figure 5. DVL2 ubiquitylation regulates interactions. **(A)** Immunofluorescence microscopy analysis was performed on MDA-MB-231 breast cancer cells expressing FLAG-DVL2. **(B)** Expanded graph of SILAC-MS analysis in **FIG 5C** showing outlier DVL2 peptides**. (C)** Description of outlier DVL2 peptides shown in **(B)** with corresponding normalized H:L ratio.

**Supplemental Figure S6.**
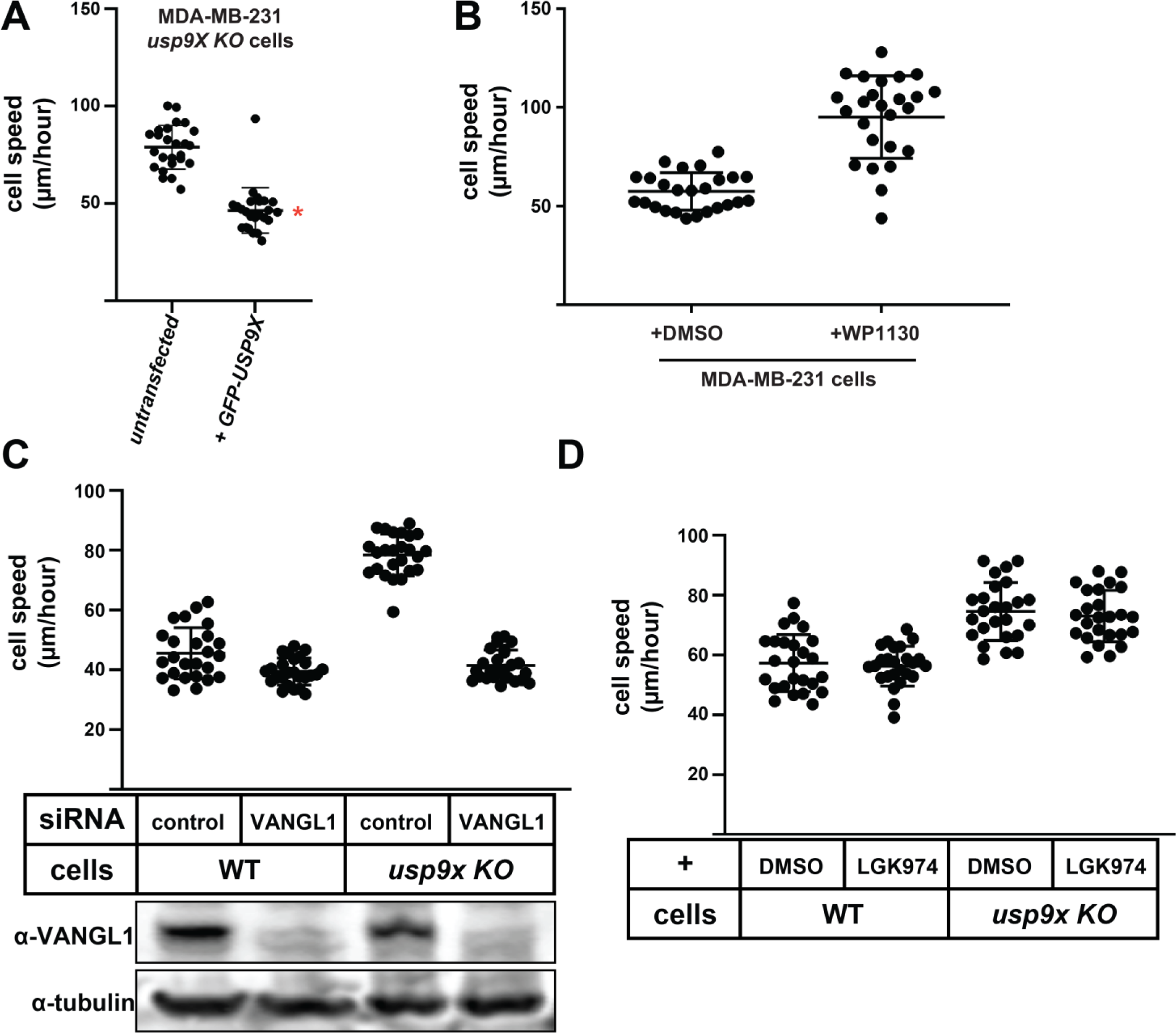
Related to Figure 6. USP9X antagonizes cell motility. **(A)** The indicated MDA-MB-231 cell lines were transiently transfected with a GFP-USP9X expression vector and cell migration speed was measured for untransfected and transfected (GFP-positive) cells (n=25). The red asterisk represents a statistically significant difference compared to untransfected *usp9x* KO cells. **(B)** Migration speed of MDA-MB-231 cells was measured after treatment with DMSO control or WP1130 (300nM). There is a statistically significant difference when comparing cell speed after WP1130 treatment to DMSO-treated control (p<0.005). **(C)** Migration speed of MDA-MB-231 cells was measured after treating with control siRNA or siRNA targeting VANGL1. There was a statistically significant difference (p<0.05) in cell speed with VANGL1 knockdown when compared to control siRNA treatment in the indicated cell lines. Bottom panel: Corresponding immunoblots confirming VANGL1 knockdown with siRNA treatment. **(D)** Migration speed of indicated MDA-MB-231 cells after treatment with DMSO control or LGK974 (1nM). There was no significant difference with LGK974 treatment when compared to DMSO control in indicated cell types.

**Supplemental Figure 7.**
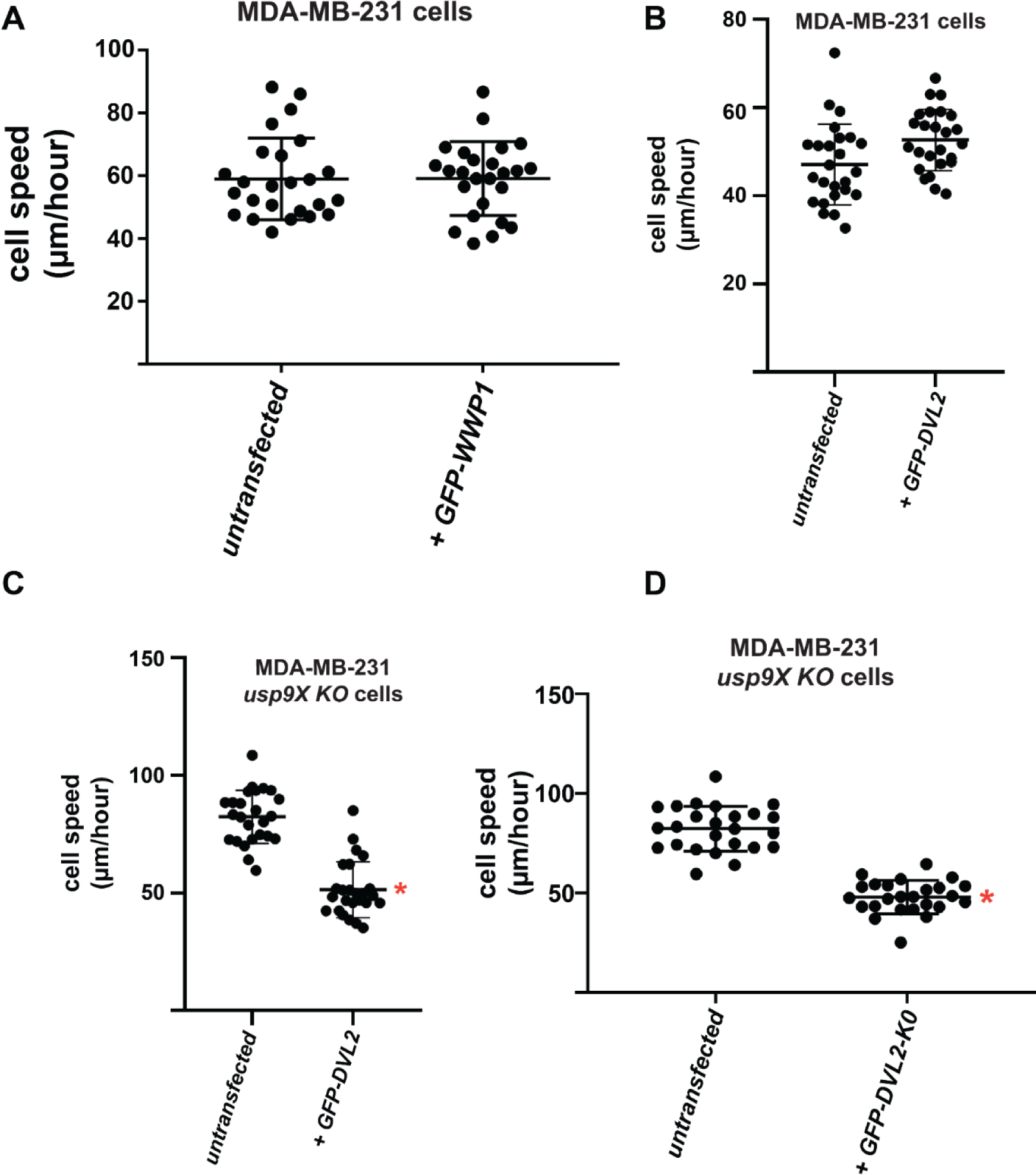
Related to Figure 7. DVL2 expression suppresses the motility phenotype of *usp9x* knockout cells. **(A)** The indicated MDA-MB-231 cell lines were transiently transfected with a GFP-WWP1 expression vector and cell migration speed was measured for untransfected and transfected (GFP-positive) cells (n=25). There was no significant difference relative to untransfected cells. **(B-C)** The indicated MDA-MB-231 cell lines were transiently transfected with a GFP-DVL2 expression vector and cell migration speed was measured for untransfected and transfected (GFP-positive) cells (n=25). Red asterisk indicates a statistically significant difference (p<0.005) when compared to untransfected control in the indicated cell line (n=25). **(D)** The indicated MDA-MB-231 cell lines were transiently transfected with a GFP-DVL2-K0 (a construct in which all lysines have been mutated to arginines) expression vector and cell migration speed was measured for untransfected and transfected (GFP-positive) cells (n=25). Red asterisk indicates a significant difference (p<0.005) when compared to untransfected control in the indicated cell line (n=25).

**Table S1.** WWP1 Interaction Profile using SILAC-MS

| category | protein | known function / description | pept. count | H:L ratio | stoich. (norm.) |
| --- | --- | --- | --- | --- | --- |
| **Bait** | WWP1 | NEDD4 family E3 ubiquitin ligase (bait) | 289 | 36.01 | 1.00 |
| **membrane traffic** | SCAMP3 | endosomal recycling | 18 | 37.43 | 0.16 |
|  | ARF4 | polarized exocytosis | 8 | 7.13 | 0.14 |
|  | RAB14 | regulation of Golgi-to-endosome transport | 5 | 9.22 | 0.07 |
|  | ARF5 | endocytosis of integrins | 3 | 12.40 | 0.05 |
|  | SEC23IP | regulates ER exit | 9 | 5.47 | 0.03 |
| **plasma membrane / membrane associated** | CCDC85C | function unknown | 21 | 17.47 | 0.16 |
|  | SPG20 | function unknown | 33 | 47.71 | 0.16 |
|  | AMOTL2 | regulation of cell junctions; Hippo signaling | 22 | 38.39 | 0.09 |
|  | AMOTL1 | regulation of cell junctions; Hippo signaling | 21 | 33.19 | 0.07 |
|  | RAPGEF6 | function unknown | 31 | 28.00 | 0.06 |
|  | MC1R | melanocortin receptor | 3 | 26.71 | 0.03 |
|  | AXL | receptor tyrosine kinase | 8 | 5.18 | 0.03 |
|  | ERBB2IP | binds and stabilizes ERBB2 receptor | 12 | 9.36 | 0.03 |
|  | FNDC3B | function unknown | 10 | 5.19 | 0.03 |
|  | SLC2A1 | facilitative glucose transporter | 3 | 6.67 | 0.02 |
|  | LRP10 | putative receptor; function unknown | 4 | 12.62 | 0.02 |
|  | EGFR | epidermal growth factor receptor | 6 | 8.66 | 0.02 |
|  | ADAM9 | cell surface metalloproteinase | 4 | 6.04 | 0.02 |
|  | NPC1 | intracellular cholesterol transporter | 4 | 8.53 | 0.01 |
| **signaling** | WBP2 | Hippo pathway and hormone signaling | 26 | 17.07 | 0.32 |
|  | PDLIM7 | bone formation and BMP signaling pathways | 30 | 22.18 | 0.21 |
|  | PTPN14 | protein tyrosine phosphatase | 75 | 22.11 | 0.20 |
|  | TMEM55B | phosphatidylinositol 4,5-bisphosphate 4-phosphatase | 4 | 6.14 | 0.05 |
|  | DVL2 | signal transducer, WNT pathway | 9 | 25.67 | 0.04 |
|  | PTPN21 | protein tyrosine phosphatase | 4 | 10.24 | 0.01 |
| **ubiquitin proteasome system** | USP9X | deubiquitylase | 94 | 25.37 | 0.12 |
|  | FAM175B | member of the BRISC complex; deubiquitylase | 9 | 13.22 | 0.07 |
|  | TXNIP | thioredoxin interacting protein; may function as E3 adaptor | 8 | 8.86 | 0.06 |
|  | BRCC3 | member of the BRISC complex; deubiquitylase | 5 | 21.72 | 0.05 |
|  | ARRDC1 | thioredoxin interacting protein; may function as E3 adaptor | 6 | 20.14 | 0.04 |
|  | C19ORF62 | member of the BRISC complex; deubiquitylase | 3 | 9.28 | 0.03 |
|  | USP24 | deubiquitylase | 20 | 23.66 | 0.02 |

**Light:** MDA-MB-231 cells stably expressing pQCXIP (empty vector)
**Heavy:** MDA-MB-231 cells stably expressing FLAG-WWP1
**Selection Criteria:** (1) number of peptides quantified $\geq 3$ ; (2) H:L ratio $> 5$

**Table S2.**
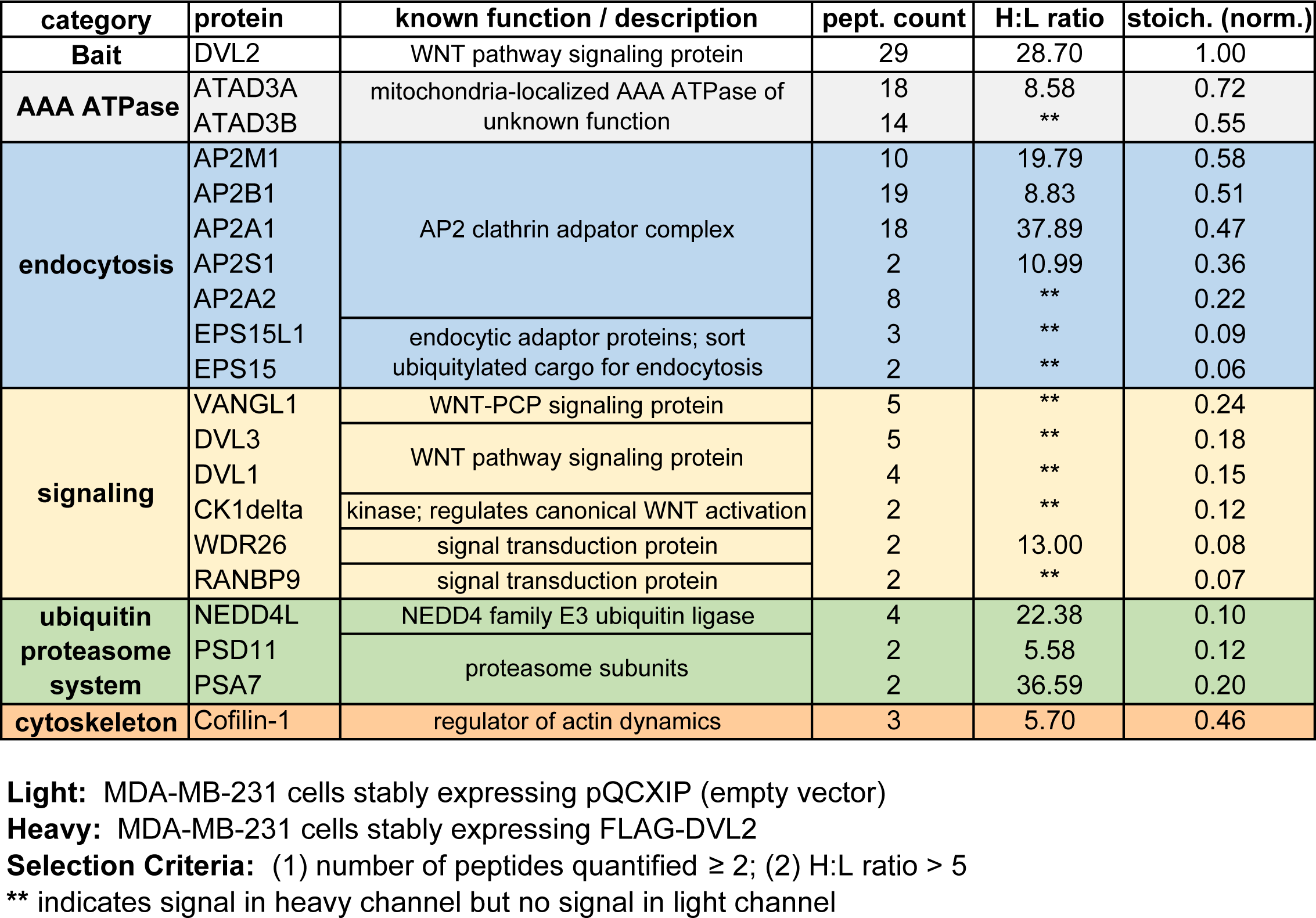
DVL2 Interaction Profile using SILAC-MS

**Table S3.** DVL2 Interaction Profile in WNT-activated cells using SILAC-MS

| category | Protein | known function / description | pept. count | H:L ratio | stoich. (norm.) |
| --- | --- | --- | --- | --- | --- |
| bait | DVL2 | WNT pathway signaling protein | 43 | 107.20 | 1.00 |
| AAA ATPase | ATAD3B | mitochondria-localized AAA ATPase of unknown function | 15 | ** | 0.40 |
|  | ATAD3A |  | 25 | 62.37 | 0.67 |
| endocytosis and trafficking | AP2B1 | AP2 clathrin adaptor complex | 49 | 52.27 | 0.90 |
|  | AP2M1 |  | 17 | 43.41 | 0.67 |
|  | AP2A1 |  | 38 | 60.82 | 0.67 |
|  | AP2S1 |  | 5 | 71.38 | 0.60 |
|  | AP2A2 |  | 30 | 27.94 | 0.55 |
|  | CAVIN1 | caveolae formation and organization | 10 | 18.36 | 0.44 |
|  | CAVIN2 |  | 8 | ** | 0.32 |
|  | SQSTM1 | autophagy receptor | 10 | 10.15 | 0.39 |
|  | EPS15L1 | endocytic adaptor proteins; sort ubiquitylated cargo for endocytosis | 7 | 6.31 | 0.14 |
|  | EPS15 |  | 2 | ** | 0.04 |
| signaling | VANGL1 | WNT-PCP signaling protein | 13 | ** | 0.42 |
|  | CK1 delta | casein kinase isoforms; regulate canonical WNT activation | 17 | ** | 0.70 |
|  | CKII beta |  | 7 | ** | 0.56 |
|  | CK1 epsilon |  | 12 | ** | 0.49 |
|  | CKII alpha |  | 9 | ** | 0.39 |
|  | PLK1 | polo-like kinase-1 | 16 | ** | 0.45 |
|  | DVL3 | WNT pathway signaling protein | 15 | ** | 0.36 |
|  | RANBP9 | signal transduction protein | 15 | ** | 0.35 |
|  | WDR26 | signal transduction protein | 13 | 74.44 | 0.34 |
| ubiquitin proteasome system | NEDD4L | NEDD4 family E3 ubiquitin ligase | 29 | 35.38 | 0.51 |
|  | WWP2 |  | 12 | ** | 0.24 |
|  | WWP1 |  | 3 | ** | 0.06 |
|  | PSA7 | proteasome subunits | 2 | ** | 0.14 |
|  | PRS10 |  | 3 | ** | 0.13 |
|  | PSA1 |  | 2 | ** | 0.13 |
|  | PSD11 |  | 2 | ** | 0.08 |
| transcription | FOXK2 | transcriptional regulators | 18 | ** | 0.47 |
|  | FOXK1 |  | 13 | ** | 0.30 |
| protein folding | BAG3 | co-factor for HSP70 | 14 | ** | 0.42 |

**Light:** MDA-MB-231 cells stably expressing pQCXIP (empty vector) stimulated with 200ng/mL mouse-Wnt3a and 100ng/mL mouse-Rspondin
**Heavy:** MDA-MB-231 cells stably expressing FLAG-DVL2 stimulated with 200ng/mL mouse-Wnt3a and 100ng/mL mouse-Rspondin
**Selection Criteria:** (1) number of peptides quantified $\geq 2$ ; (2) H:L ratio $> 5$
\*\* indicates signal in heavy channel but no signal in light channel

